# Shedding light on biosilica morphogenesis by comparative analysis of the silica-associated proteomes from three diatom species

**DOI:** 10.1101/2021.09.18.460806

**Authors:** Alastair W. Skeffington, Marc Gentzel, Andre Ohara, Alexander Milentyev, Christoph Heintze, Lorenz Böttcher, Stefan Görlich, Andrej Shevchenko, Nicole Poulsen, Nils Kröger

## Abstract

- Morphogenesis of the intricate patterns of diatom silica cell walls is a protein-guided process, yet to date only very few such silica morphogenetic proteins have been identified. Therefore, it is unknown whether all diatoms share conserved proteins of a basal silica forming machinery, and whether unique proteins are responsible for the morphogenesis of species specific silica patterns.
- To answer these questions, we extracted proteins from the silica of three diatom species (*Thalassiosira pseudonana, Thalassiosira oceanica* and *Cyclotella cryptica*) by complete demineralization of the cell walls. LC-MS/MS analysis of the extracts identified 92 proteins that we name ‘Soluble Silicome Proteins’ (SSPs).
- Surprisingly, no SSPs are common to all three species, and most SSPs showed very low similarity to one another in sequence alignments. In depth bioinformatics analyses revealed that SSPs can be grouped into distinct classes bases on short unconventional sequence motifs whose functions are yet unknown. The results from *in vivo* localization of selected SSPs indicates that proteins, which lack sequence homology but share unconventional sequence motifs may exert similar functions in the morphogenesis of the diatom silica cell wall.

## Introduction

Silica is the hydrated, amorphous oxide of the element silicon (SiO_2_·nH_2_O), and is the second most abundant biologically formed mineral (biomineral) after calcium carbonate (Lowenstam & Weiner, 1989). It occurs in all eukaryotic supergroups (Marron *et al*., 2016), but the biological production of silica is dominated by a group of micro-algae called diatoms (Nelson *et al*., 1995). Oceanic diatoms alone account for about 20% of global primary biological production (Benoiston *et al*., 2017), and therefore closely link the biogeochemical cycles of silicon and carbon. Diatoms use silica as their cell wall material, which is believed to have been a key factor in their ecological success, functioning as an armour against predators (Pančić *et al*., 2019), improving nutrient uptake (Mitchell *et al*., 2013), providing photoprotection, and as a means to regulate buoyancy (Raven & Waite, 2004). Furthermore, biogenesis of the intricately nano- and micropatterned silica cell walls of diatoms has attracted great interest from materials scientist as paradigms for the bottom-up production of 3D, hierarchically porous materials (Nassif & Livage, 2011).

Silica formation in diatoms occurs in specialized, membrane-bound, intracellular compartments called silica deposition vesicles (SDVs). There are two different types of SDVs. One produces plate- or dome-shaped silica structures called valves, and the other produces biosilica rings, called girdle bands. During cell division, each daughter cell produces one valve, while several girdle bands are produced during interphase (Hildebrand *et al*., 2007). When morphogenesis of a valve or girdle band inside an SDVs has been completed, the silica structure is exocytosed and assembled into a cell wall that completely encases the cell.

It has been proposed that the lumen of each SDV possesses a silica-forming matrix composed of organic macromolecules (Hecky *et al*., 1973; Volcani, 1981; Kröger & Sumper, 2004). In this model, the species-specific silica nano- and micropatterns would result from differences in the composition of the organic matrix components. Currently, the biomolecular composition of SDVs is unknown, because a method for their isolation has not yet been established. However, it is assumed that some of the organic molecules associated with the mature cell wall originate from the silicification machinery of the SDV (Hecky *et al*., 1973). Complete demineralization of diatom cell walls using an acidic ammonium fluoride solution, solubilised phosphoproteins and long-chain polyamines (Kröger & Sumper, 2004), while organic matrices composed of proteins and polysaccharides remained insoluble (Brunner *et al*., 2009a; Scheffel *et al*., 2011; Buhmann *et al*., 2014; Kotzsch *et al*., 2017; Tesson *et al*., 2017; Pawolski *et al*., 2018). In vitro silica formation experiments using mixtures of the phosphoproteins and long-chain polyamines, both in the presence and absence of insoluble organic matrices yielded porous silica patterns that mimicked native silica morphologies, suggesting that the biosilica associated organic components might play a role in silica morphogenesis *in vivo* (Poulsen *et al*., 2003; Poulsen & Kröger, 2004; Scheffel *et al*., 2011; Pawolski *et al*., 2018).

To date, diatom biosilica associated proteins have been biochemically characterized from only two diatom species, *Cylindrotheca fusiformis* (Kröger *et al*., 2001, 2002; Poulsen *et al*., 2003) and *Thalassiosira pseudonana* (Poulsen & Kröger, 2004; Scheffel *et al*., 2011; Kotzsch *et al*., 2016). The most prominent groups of proteins, called silaffins, exhibit no significant sequence similarity to each other or to any other proteins in the NCBI database, but they share a very high degree of phosphorylation and unusual polyamine modifications of lysine residues (Poulsen & Kröger, 2004; Scheffel *et al*., 2011). A large number of genes have been identified that are responsive to silicic acid content in the culture medium (Mock *et al*., 2008; Sapriel *et al*., 2009) or upregulated during valve formation (Shrestha *et al*., 2012; Brembu *et al*., 2017). However, for almost all of these genes it is unknown whether the encoded proteins are associated with the biosilica or involved in silica morphogenesis.

In the present study we aimed to substantially expand our knowledge of the silica associated proteome of diatoms, and to identify proteins that might be responsible for differences in silica morphology between species. To achieve this, we chose three closely related species of centric diatoms: *T. pseudonana, Thalassiosira oceanica* and *Cyclotella cryptica*. It should be noted that the genus Thalassiosira is paraphyletic (Kaczmarska *et al*., 2005), and *T. pseudonana* is more closely related to *C. cryptica* than it is to *T. oceanica* (Alverson *et al*., 2011). These species have a conserved architecture of cylindrically shaped cell walls, with a complex silica network structure in the valves. The network consists of radially oriented ribs, which form a branched pattern from the centre of the valve to its rim (Fig. **1a, g, b, h** yellow and green lines), and which are more pronounced in *C. cryptica* than in *T. pseudonana*. Compared to the other two species, *T. oceanica* differs in that the ribs are obscured by a covering layer of porous silica (Fig. **1d, e**). Neighbouring ribs are connected by seemingly irregularly spaced short silica bridges in *C. cryptica* and *T. pseudonana*. The gaps in the meshwork of interconnected ribs and silica bridges contain silica that is perforated with 20 – 30 nm sized circular pores (known as cribrum pores; Fig. **1b, h**). Multiple tube-like structures, termed fultoportulae, are regularly spaced near the rim of the valves in all three species (Fig. **1a, d, g, b, e, h**, red circles). About half of the valves also contain one or two fultoportulae near the centre (Fig. **1a, d**, purple circle). Fultoportulae are involved in the secretion of chitin fibres (Herth, 1979) and have a restricted phylogenetic distribution within the centric diatoms (Kaczmarska *et al*., 2005). The girdle band regions of the cell walls are less complex than the valves, exhibiting a pattern of alternating porous and non-porous regions (Fig. **1c, f, i**).

**Figure 1.**
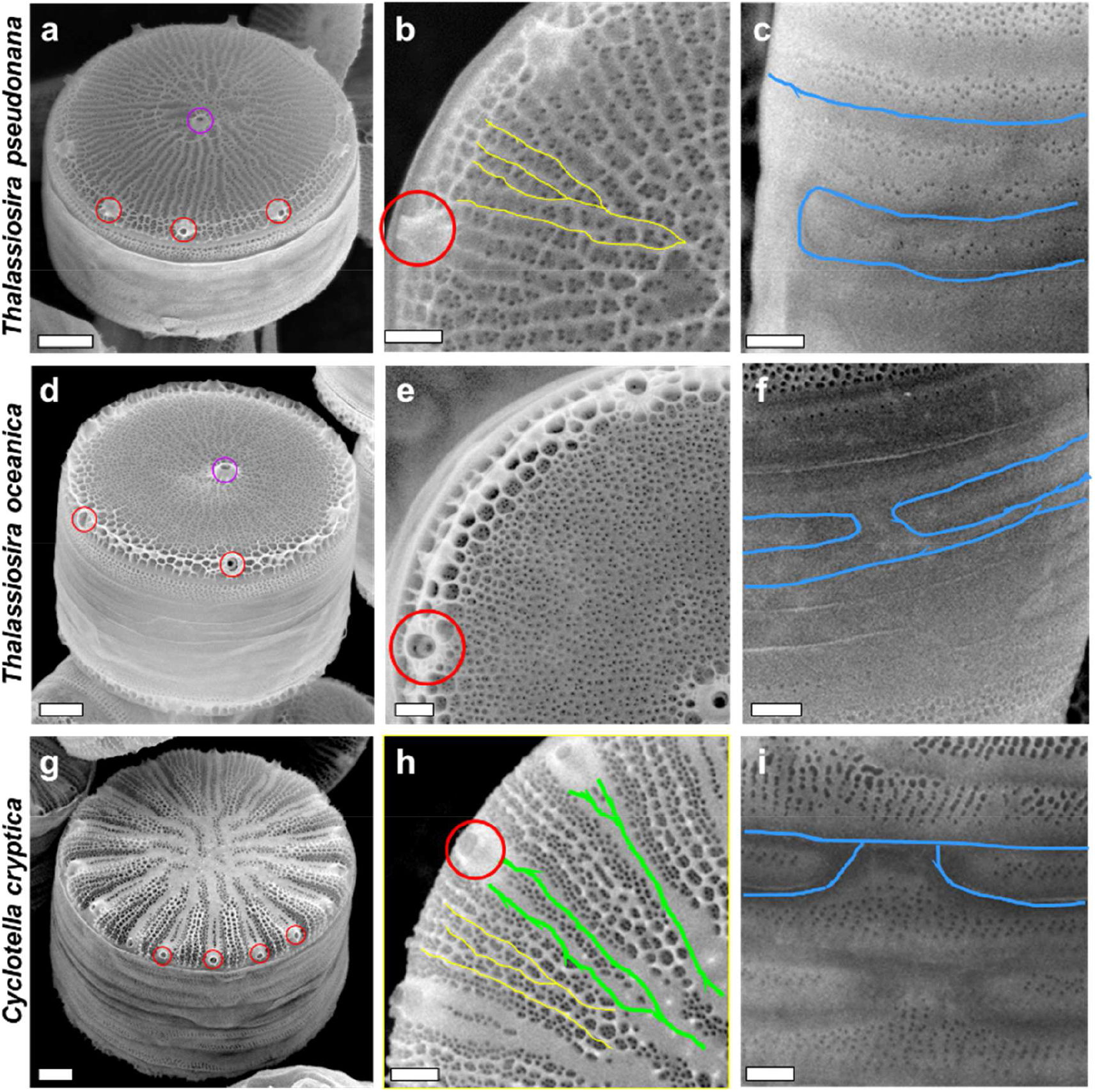
Scanning electron microscopy (SEM) images of isolated silica cell walls from *T. pseudonana* (a – c), *T. oceanica* (d – f) and *C. cryptica* (g – i). For each species, the cell wall is presented in oblique view (a, d, g), and details from the valve region (b, e, h) and girdle band region (c, f, i) are shown. Yellow and green lines indicate radial ribs and wide radial ribs, respectively. Blue lines designate non porous regions in girdle bands. Examples of fultoportulae on the rim are labeled with red circles, and central fultoportulae with purple circles. Scale bars: 1 µm (a, d, g) and 400 nm (b, c, e, f, h, i).

In this work we identify soluble silica associated proteins from these three species, thereby shedding light on the machinery that might be responsible for the commonalities and differences in silica structure, as well as identifying more conserved proteins that might have a more basal role in silica biogenesis.

## Materials and Methods

### Culture conditions

*Thalassiosira pseudonana* (Hustedt) Hasle et Heimdal clone CCMP1335, *Cyclotella cryptica* Reimann, Lewin et Guillard strain CCMP332 and *Thalassiosira oceanica* (Hustedt) Hasle et Heimdal CCMP1005 were grown in an enriched artificial seawater medium (EASW) (Harrison *et al*., 1980) at 18°C under constant light at 5,000-10,000 lux.

### Identification of silica cell wall associated proteins

Isolation of diatom silica and subsequent ammonium fluoride extraction was performed as described for *T. pseudonana* (Kotzsch *et al*., 2016). The ammonium fluoride extracts were centrifuged at 3200 g for 30 min and the supernatants were desalted by three rounds of ultrafiltration (MWCO: 3-6 kDa), each resulting in a 10-fold dilution with 10 mM ammonium acetate. Extracts were analysed by SDS-PAGE using 6% Tris-Tricine gels (Schägger & von Jagow, 1987) stained using “Stains-All” (Campbell *et al*., 1983). In solution and in-gel digests were analysed by LC-MS/MS and database searches carried out using Mascot (Methods **S1**) (Matrixscience, UK). Protein inference was carried out using Scaffold (ProteomeSoftware, USA) (protein probability > 0.99, at least two unique peptides, found in two biological replicates: see Methods **S1**). Raw data have been deposited in the PRIDE database (Chambers *et al*., 2012) (part of the ProteomeXchange consortium (Deutsch *et al*., 2020)), with accession PDX026496.

### Bioinformatic methods

The properties of the protein sequences were investigated using ProminTools (Skeffington & Donath, 2020), which relies on various software tools: low complexity regions were identified using Seg (Wootton & Federhen, 1993) with default parameters; predicted intrinsic disorder was calculated using VSL2 (Peng *et al*., 2006); biases in amino acid content were identified using the fLPS software (Harrison, 2017) with a p-value cut-off of 10^−6^ and bias quantified as described in Skeffington and Donnath, 2020. Motif finding was carried out using Motif-x (Chou & Schwartz, 2011) via ProteinMotifFinder (Skeffington & Donath, 2020). Clustering based on motif content was based on the Ward.D method, and a distance matrix based on the Distance Correlation (Székely & Rizzo, 2014) measure (further details in Methods **S2**). Pairwise global sequence alignments were calculated using EMBOSS Needle (Needleman & Wunsch, 1970) with default parameters. Cellular localisations were predicted using Hectare (Gschloessl *et al*., 2008) and ASAfind (Gruber *et al*., 2015), while transmembrane prediction was carried out using PureseqTM (Qing *et al*., 2019) and MEMSAT-SVM (Nugent & Jones, 2009). Known domains were discovered using InterPro v. 84.0 (Jones *et al*., 2014).

The phylogenetic distribution of proteins was carried out using the MMETSP transcriptome data (Keeling *et al*., 2014) in particular the “_clean.fasta” files from which contaminants have been removed (software: https://github.com/kolecko007/mmetsp_cleanup, data: www.imicrobe.us/#/projects/104) (Marron *et al*., 2016). tBLASTn searches were carried out with Seg based filtering of low complexity regions turned on, with default parameters. For further computational details see Methods **S3**.

### Generation and characterisation of transgenic cell lines

The construction for gfp fusions with genes of interest are described in detail in Methods **S4**. Introduction of fusion genes into *T. pseudonana* and *C. cryptica* and selection of transformants were performed as described previously (Poulsen *et al*., 2006; Kumari *et al*., 2020). Cells originating from least 10 independent colonies were examined by epifluorescence microscopy to check for a consistent location phenotype for each GFP fusion protein. Confocal microscopy was performed with live cells and extracted silica from representative cell lines (see Methods **S5**).

## Results

### Extraction of silica associated proteins and protein identification

Biosilica was isolated from *T. pseudonana, T. oceanica* and *C. cryptica* cultures using a previously established method for diatoms (Kröger et al., 2000), followed by complete demineralization with ammonium fluoride at pH 4.5 (Kotzsch *et al*., 2016). The soluble extracts were separated by SDS-PAGE and stained with “Stains-All”, which revealed multiple bands in the extract of *T. pseudonana* and *C. cryptica* with apparent molecular masses ranging from ∼15 kDa to >170 kDa (Fig. 2). In contrast, the *T. oceanica* extract was dominated by a broad band ranging from ∼55 kDa to ∼130 kDa. Two replicate ammonium fluoride soluble extracts for each of the three species were subjected to proteolytic digestion and subsequently analysed by liquid chromatography coupled tandem mass spectrometry (Methods **S1**). After protein identification and inference (see Methods **S1**), we discarded seven proteins as potential contaminants. Four of these were proteins with a well predicted chloroplast transit peptide (‘high confidence’ ASAfind predictions (Gruber et al., 2015)), two were identified as histones, and one as a conserved snoRNA binding protein. The remaining 92 proteins were taken forward for downstream analyses (Supporting Information Table **S1**): 29 proteins from *C. cryptica*, 55 proteins from *T. pseudonana*, and eight proteins *from T. oceanica*. These proteins were collectively termed “Soluble Silicome Proteins” (SSPs).

**Fig. 2.**
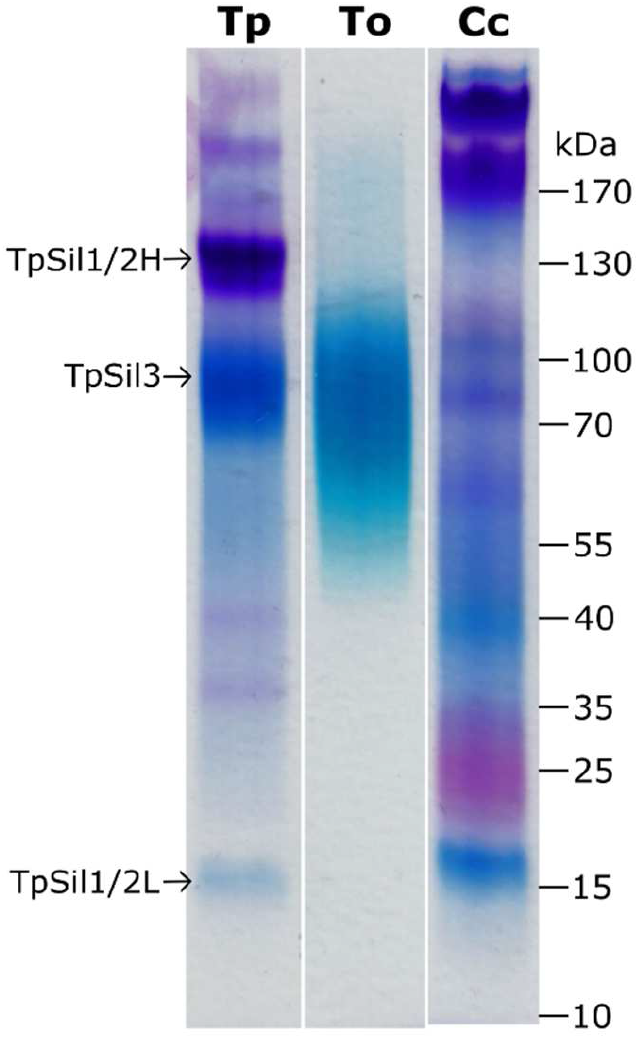
SDS-PAGE analysis of the ammonium fluoride soluble extracts. The samples were run on the same gel and were stained with the dye “Stains-All”. Tp: *T. pseudonana*, To: *T. oceanica*, Cc: *C. cryptica*. Arrows indicate previously characterized *T. pseudonana* silaffins (Poulsen & Kröger, 2004).

We considered the possibility that the lower complexity of the *T. oceanica* samples could be an artefact due to a poor *T. oceanica* database, resulting in proteins not being identified that are in fact present in all three species in reality. To examine this possibility, we performed searches of the *T. oceanica* derived spectra against the *T. pseudonana* and *C. cryptica* databases. These searches did not result in any hits, supporting the SDS-PAGE data in indicating that the lower complexity is a genuine feature of the *T. oceanica* samples.

### Previously characterised proteins and known domains

Seven of the 92 SSPs, all from *T. pseudonana*, had previously been identified at the protein level: Tp_p150 (Davis *et al*., 2005), TpSil1 and TpSil3 (Poulsen & Kröger, 2004), TpSil4 (Sumper & Brunner, 2008), SiMat3, SiMat4 and SiMat7 (Silacanin-1, Sin1) (Kotzsch et al. 2016). The 85 novel proteins were distinguished by an acronym for the species of origin and a numeric identifier (e.g. TpSSP1). A role of the SSPs in silica formation was supported by the fact that 35 of the *T. pseudonana* SSPs had been previously identified in transcriptomic experiments relating to silica metabolism (Table **S2**) (Mock *et al*., 2008; Shrestha *et al*., 2012; Brembu *et al*., 2017), and 13 were identified in more than one of the previous studies.

Twenty-nine of the SSPs contained known protein domains. Chitin recognition domains were found in three proteins (CcSSP6, CcSSP10 and Tp_p150), which is not unexpected as chitin has been found associated with diatom silica (Durkin *et al*., 2009; Brunner *et al*., 2009b). Two proteins (TpSSP7, TpSSP35) contained serine protease domains, and TpSSP44 is a predicted glycosyltransferase (family 8). These three enzymes might be involved in the post-translational processing of silaffins, which requires O-glycosylation, and multiple, site-specific proteolytic cleavage steps (Kröger *et al*., 1999; Poulsen *et al*., 2003; Poulsen & Kröger, 2004). TpSSP37 contains a DHH phosphatase domain and CcSSP26 and TpSSP45 have kinase domains, indicating that these proteins may be involved in regulating the phosphorylation patterns of silica associated phosphoproteins such as silaffins and silacidins (Kröger *et al*., 2002; Poulsen & Kröger, 2004; Wenzl *et al*., 2008). Certain silaffins have also been shown to be sulfated (Poulsen *et al*., 2003; Poulsen & Kröger, 2004), so the sulfatase CcSSP29 and the sulfotransferase CcSSP22 could potentially be involved in regulating patterns of silaffin sulfation. CcSSP21 contains a peroxidase domain, which might play a role in formation of the biosilica-associated insoluble organic matrices that have been identified in *C. cryptica, T. pseudonana* and other diatoms (Scheffel *et al*., 2011; Tesson *et al*., 2013; Pawolski et al., 2018). Although the chemical mechanisms for the formation of the insoluble organic matrices is yet unknown, it might involve H_2_O_2_-mediated cross-linking of tyrosine-bearing proteins analogous to extracellular matrix formation in the green alga *Chlamydomonas reinhardtii* (Waffenschmidt *et al*. 1993).

An alkaline phosphatase (AP) domain was found in TpSSP1, and is unlikely to be involved in silica formation, especially since the acidic pH in the SDV lumen (Vrieling *et al*., 1999) is incompatible with AP activity (Thompson & MacLeod, 1974). APs are abundant secretory proteins in diatoms and required for the acquisition of inorganic phosphate (Fuentes *et al*., 2014). It is conceivable that some APs remain stably bound to the cell wall to enhance AP activity near the cell surface. CcSSP16 is a phytase-like esterase which could also have a role in phosphate acquisition. Alternatively, it could play a role in the catalysis of silicic acid condensation, in a manner analogous to the hydrolase Silicatein (Cha *et al*., 1999).

CcSSP1 contains a fasciclin domain, and was identified with substantially more spectral counts than any other *C. cryptica* SSP (Table **S1**). Fasciclin domains are ancient structural motifs found in extracellular proteins present in all kingdoms of life (Seifert, 2018). There is precedence for fasciclin domains in unicellular algae, since they are present in proteins of the insoluble hydroxproline-rich glycoprotein framework of the Chlamydomonas cell wall (Voigt *et al*., 2010), and were identified in some putative adhesive proteins from diatoms *Phaeodactylum tricornutum* and *Amphora coffeaeformis* (Willis *et al*., 2014; Buhmann *et al*., 2016; Lachnit *et al*., 2019). Some fasciclin domain bearing proteins have functions unrelated to cell-cell or cell-surface adhesion, including secondary metabolism (Thiele *et al*., 2020), and thus the role of this domain in the diatom biosilica remains open. Another cell-adhesion related domain was identified in TpSSP24, TpSSP14 and TpSSP34, all of which carry FG-GAP domains that are also present in integrins (i.e. metazoan cell-matrix adhesion receptors (Kechagia *et al*., 2019)). Six proteins were found to contain Regulator of Chromosome Condensation (RCC1) repeat domains TpSSP6, TpSSP22, TpSSP26, TpSSP30, TpSSP40, TpSSP46), two proteins contain a FMN reductase domain (TpSSP32, CcSSP18), and one protein each contains a manose-6-phosphate receptor family domain (TpSiMat4), a LisH domain (TpSSP32), an S1 domain TpSSP23), a RAP domain (TpSSP45), and Leucine rich repeats (TpSSP39). The functional significance of any of these domains in diatom biosilica remains unclear.

Eighteen proteins were predicted to have a single transmembrane domain. There is precedence for transmembrane proteins being found associated with the diatom silica since the biolsilica-assicted protein Sin1 (Kotzsch *et al*., 2017), SAP1, and SAP3 (Tesson *et al*., 2017) are type-I transmembrane proteins. These proteins are located in the SDV membrane during silica biogenesis and their membrane anchors are cleaved off during maturation of the proteins (Kotzsch *et al*. 2017; Tesson *et al*. 2017).

### Sequence characteristics of SSPs

Global sequence analysis of the SSPs revealed that they are significantly lower in sequence complexity (Fig. **3a**) than other proteins from the respective proteomes of the three species (hereafter termed the ‘background proteomes’: predicted proteome minus the SSPs). They are also predicted to be disordered over a significantly greater proportion of their length than the background proteomes (Fig **3b**), and display substantial biases in amino acid composition (Fig. **S1**). The probability of achieving bias at least this extreme through random sampling of the proteome is less than 0.001 for *T. pseudonana* and *C. cryptica*, and less than 0.01 for *T. oceanica*.

**Fig. 3.**
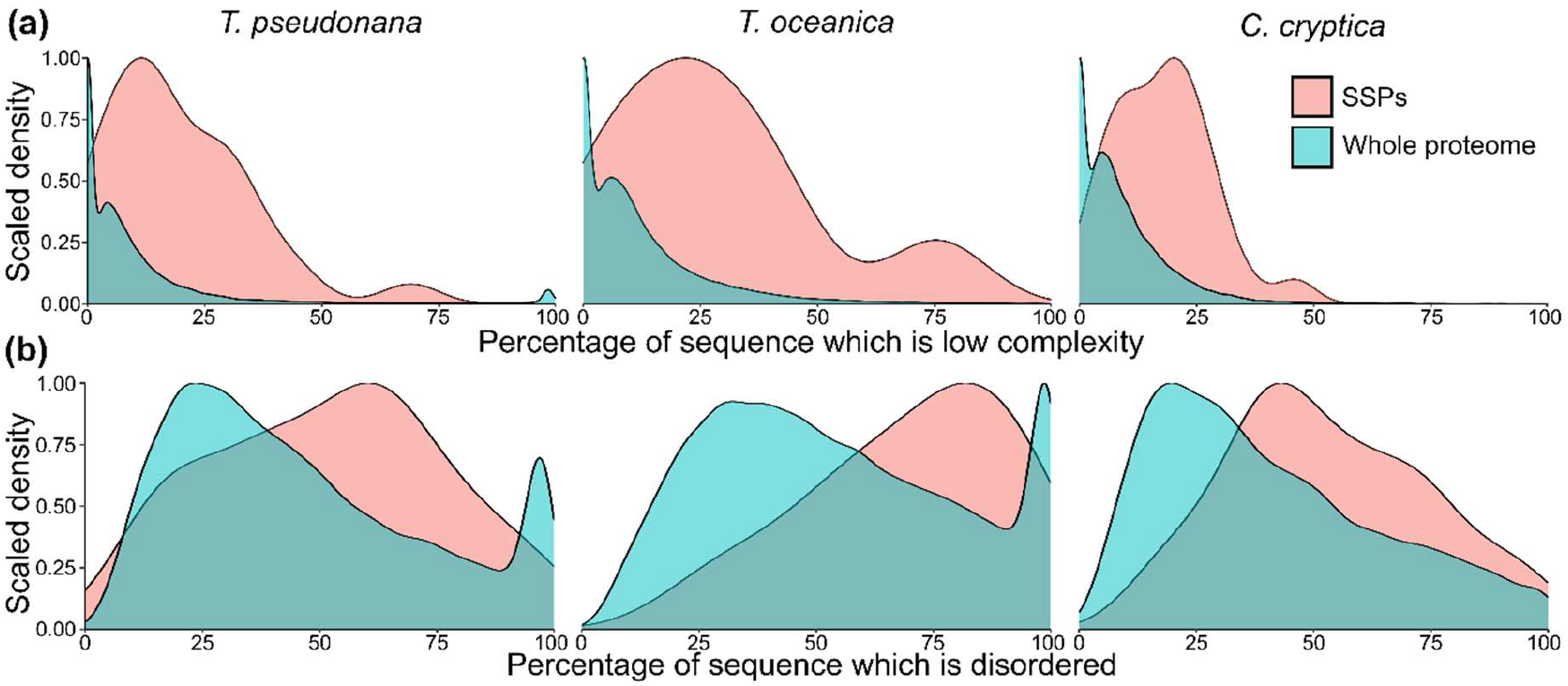
Global properties of the SSPs from *C. cryptica*, and *T. oceanica*, and *T. pseudonana*. Density plots of the percentage of (**a**) low complexity sequence in individual proteins and (**b**) predicted disordered sequence in individual proteins. In each case, the SSPs and background proteome distributions are significantly different (Wilcoxon rank sum test with continuity correction, *p* < 10^−6^).

Sequence similarity among SSPs is generally low, and no protein has putative homologues in the SSP set of all three species. In the whole dataset, there are only ten pairs of proteins with an identity greater than 35% in global sequence alignments (Table **S3**). Several groups of these proteins included both *T. pseudonana* and *C. cryptica* proteins, suggesting that there is some conservation in the silica associated proteomes of these species. For example, *C. cryptica* homologues of TpSil1 and Tp_p150 were identified as CcSSP6 and CcSSP10 respectively. TpSSP2 and CcSSP2 are also noteworthy since they are 44% identical and were both identified with high (normalized) spectral counts.

There are no proteins with good similarity (BLAST e-value < 1e-30 and query coverage > 50%) to the *T. oceanica* SSPs represented in either the proteomes or the genomes of the other two species.However the *T. pseudonana* and C. cryptica SSPs did have similar proteins in the *T. oceanica* proteome in 55% (30 proteins) and 34% (10 proteins) respectively, although none of these were found in *T. oceanica* SSPs. Where *T. pseudonana* lacked a similar SSP to a *C. cryptica* SSP (or vice versa, based on >35% identity in global sequence alignments, 66 proteins in total) the SSPs were ‘missing’ for two different reasons. In 51% of these cases (34 proteins) there is no similar protein in the other predicted proteome (BLAST e-value < 1e-30 and qcov > 50%). In the other 49% of cases (32 proteins), a similar protein is represented in the other predicted proteome, suggesting that the protein was either not expressed at the time of cell harvesting, or failed to be detected in the proteomic analysis.

Despite the low level of sequence similarity among the SSPs at the level of whole protein sequence alignments, manual inspection of the sequences revealed that short sequence motifs were often shared between subsets of the proteins. To understand the nature of such smaller scale sequence similarities, we searched for sequence motifs that were overrepresented in the SSPs relative to the background proteomes. Enriched motifs were identified using the motif-x software (Chou & Schwartz, 2011) via the ProminTools package (Skeffington & Donath, 2020). In total, 113 motifs were found to be enriched in the SSPs relative to the background proteome (motif-x p-value cut-off = 10^−6^), reflecting the highly biased composition of the sequences (Table **S4**). None of the enriched motifs are common to all SSPs.

Previously, K..K motifs (period = any amino acid) have been identified in many silica associated proteins, and experimentally demonstrated to be involved in targeting these proteins to the cell wall (Poulsen *et al*., 2013), and to promote silica formation *in vitro* (Wieneke *et al*., 2011). The motif K..K is common in all three background proteomes and is only three-fold enriched in the SSPs. However, specific variants of K..K with restricted identities for both the central and the flanking amino acids were among the most prominent motifs in the data set. In particular, PKA.K, KS.KS, GKS.K, SKS.K, SKA.K, AKS.K, KS.KA and GKA.K were all more than 20-fold enriched. This included DAKA.K, which is 124-fold enriched and found in 8 SSPs, KSSKA is 77-fold enriched and present in 11 SSPs, and KAEK is 33-fold enriched and found in 9 SSPs.

Given that most SSPs could not be grouped based on full-length sequence comparisons, we investigated whether the proteins could be grouped based on their motif content. Distance correlation coefficients (Székely & Rizzo, 2014) were calculated for all pairs of the 92 SSPs based on their enrichment in the identified motifs (see Materials and Methods, Table **S5**). The 52 protein sequences with strong motif-based similarity to at least one other protein (dcor > 0.65) were clustered based on their enrichment in the 70 most enriched motifs. The remaining motifs tended to be enriched to a low level in a high proportion of the SSPs, and thus did not aid in grouping the sequences. This resulted in eight protein clusters (Fig. **4**), with distinct patterns of motif enrichment. Cluster 1 (10 proteins) and Cluster 4 (10 proteins) were particularly enriched in different variants of K..K motifs (especially KSGK and FKS.K in cluster one, and KSSKA in Cluster 4). Cluster 2 (three proteins) is rich in SMSM, as were a number of proteins in Cluster 5, which were additionally highly enriched in DAKA.K. Cluster 3 (four proteins) is particularly enriched in S..SSKS, while Cluster 6 (six proteins including TpSil1 and TpSiMat4) were enriched in various P, T and S containing motifs. Finally, Cluster 7 is enriched in a range of C and G containing motifs, and Cluster 8 in various N and Q containing motifs.

**Fig. 4.**
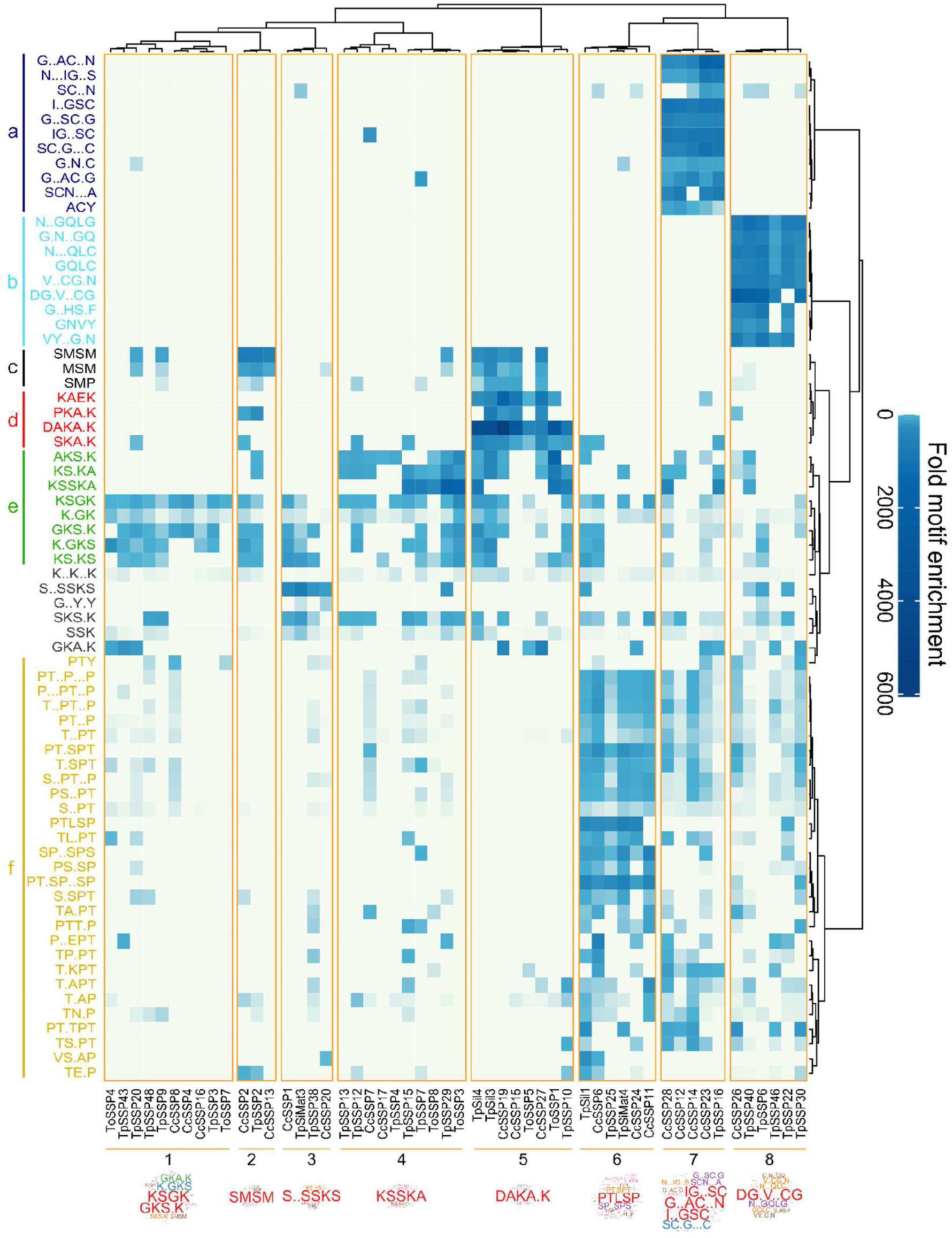
Heatmap displaying motif enrichment with respect to the relevant background proteomes. To be included in this analysis, each protein had to be highly correlated in its motif profile to at least one other protein (see Materials and Methods). Eight protein clusters were identified and are indicated at the bottom of the heatmap. For each cluster, the motif profile is represented as a wordcloud. The height of the letters is proportional to the mean enrichment of the motif in that protein cluster, but the scale differs for each wordcloud. On the left margin groups of motifs of interest (selected by co-occurrence in similar proteins and/or by similarity in amino acid composition) are coloured and labelled a – e. These correspond to the motif groups represented in Fig. 6.

Pairwise BLASTp comparisons of the proteins within the clusters (Fig. **S2**) demonstrated that the motif-based clustering revealed similarities between the proteins not captured by BLAST comparisons. In particular, in clusters one to six, only eight of the total 120 pairwise comparisons displayed BLAST similarity at an e-value cut-off of 0.01.

### Phylogenetic distribution of SSPs

Two approaches were used to assess the phylogenetic distribution of the SSPs. The first makes use of the precalculated orthogroups available at PLAZA diatoms (Osuna-Cruz *et al*., 2020), which include sequences from 10 diatom species with sequenced genomes as well as sixteen other eukaryotes. Proteins can then be classified as being species-specific or restricted to the centric diatoms, diatoms, Chromista, or as having a broad distribution across the eukaryotes. The second method was to run tBLASTn searches of the SSP sequences against transcriptomes from the Marine Microbial Eukaryote Sequencing Project (Keeling *et al*., 2014), including 136 diatom transcriptomes and 390 from other taxa. A version of the MMETSP data was used that had been cleaned of contaminants (see Materials and Methods). Of these two methods, orthogroups provide a more rigorous assessment of shared ancestry. Here they are built on genome sequence information, so proteins will not be missing due to low or absent expression as may be the case in the MMETSP transcriptomes. However, the orthology analysis does not capture the diversity of the diatoms or the eukaryotes, which is much better represented by the MMETSP transcriptomes. Overall, it is clear that neither method provides a definitive assessment of phylogenetic distribution, but rather provide complementary information to each other. The grouping of proteins by phylogenetic distribution in Fig. **5** is based on the MMETSP BLAST searches (e-value cut-off of 10^−30^ used for group assignment), and the PLAZA classifications are also indicated.

**Fig. 5.**
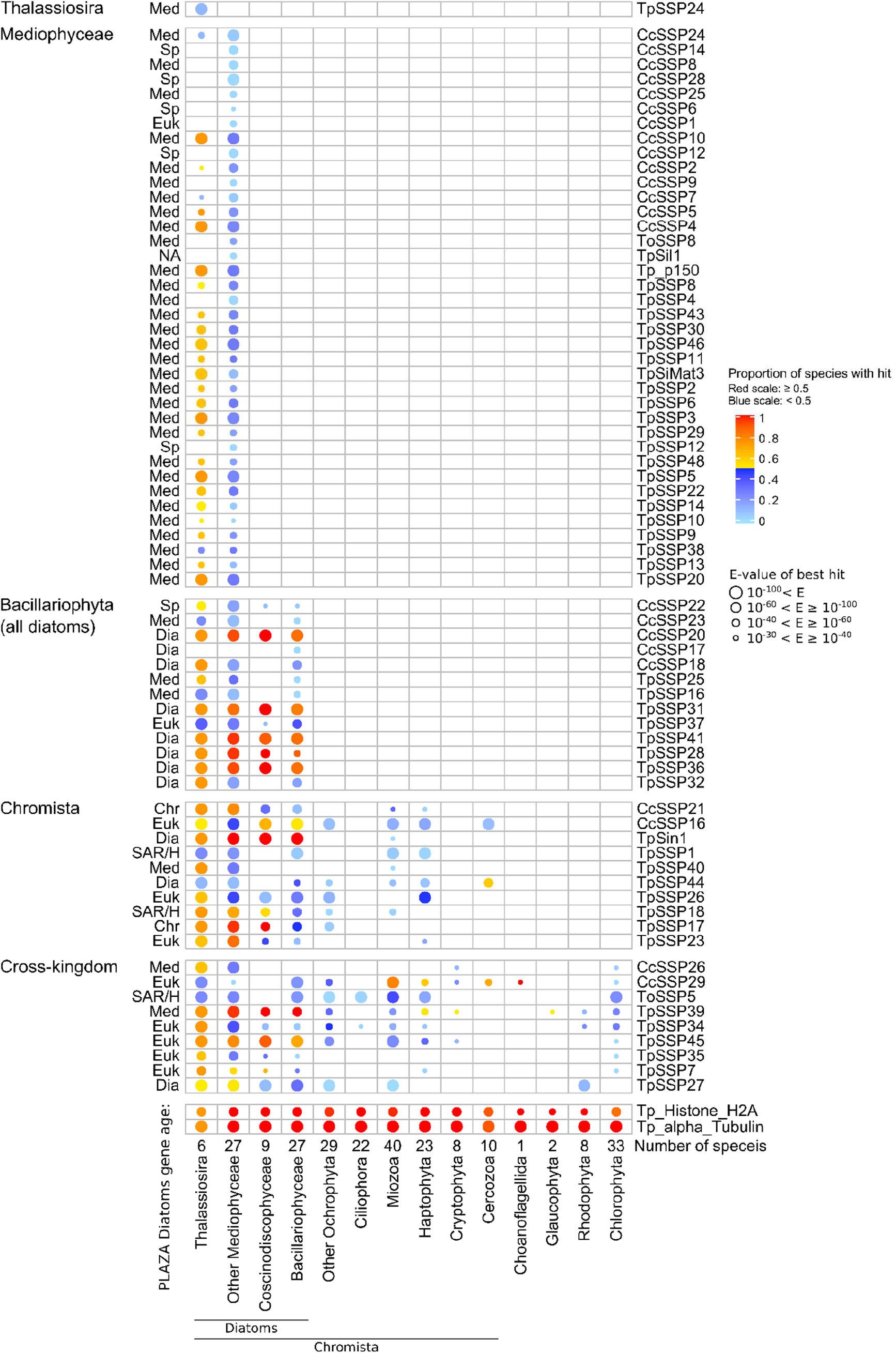
Phylogenetic distribution of SSPs. The central panel summarises the BLAST analysis of the SSPs against the MMETSP transcriptomes (see Materials and Methods. Only proteins with BLAST hits at an e-value < 10^−30^ are included in this analysis. Each column displays data from a different taxonomic group. The number of hits per species is visualised as the colour of circles, while the e-value of the best hit is indicated via the size of the circles. Proteins are classified as specific to the genus Thalassiosira, the class Mediosphyceae, the phylum Bacillariophyta (diatoms), the kingdom Chromista or as having a cross-kingdom classification. The search results for of two highly conserved proteins, α-tubulin and Histone-2A from *T. pseudonana*, are displayed for comparison. The phylogenetic distribution of the PLAZA diatom orthogroup to which each protein belongs is noted. TpSil1 was not represented in the PLAZA database (B8LDT2 is mis-annotated as TpSil1 in the database). SpSp: species specific; Med: mediophyceae specific (note PLAZA only contains Mediophyceae from the Thalassiosira clade), Dia: diatom specific, Chr: Chromista specific, Euk: Eukaryote-wide distribution.

Of the 21 proteins found to be species-specific (and therefore not displayed in Fig. **5**), nine were from *T. oceanica*, six from *C. cryptica* and six from *T. oceanica*. The largest phylogenetic grouping consisted of the 38 proteins (23 from *T. pseudonana*, 14 from *C. cryptica*, one from *T. oceanica*) restricted to the Mediophyceae (Thalassiosirales and bipolar centric diatoms). Among these, nine proteins were especially well conserved, being found in 54-67% of Mediophyceae species, which makes them good candidates for the Mediophyceae-specific silica biomineralization machinery. These proteins are: TpSiMat3; TpSSP46; Tp_p150 and its homologue CcSSP10; TpSSP3 and its homologues CcSSP4 and TpSSP5; TpSSP6 and its homologue TpSSP22.

The Bacillariophyta specific SSPs comprise 13 SSPs (eight from *T. pseudonana*, five from *C. cryptica*), which are candidates for being components of a basal, diatom specific biomineralization machinery. Of these, five proteins were identified in more than 80% of diatom species. These were TpSSP41, TpSSP28 and CcSSP20 along with its homologues TpSSP31 and TpSSP36. TpSin1 was classified as having a Chromista-wide distribution by the MMETSP analysis, due to blast hits (best hit e-value = 1.8E-36, 25% query coverage, 65% identity) in the dinoflagellates *Glenodinium foliaceum* and *Kryptoperidinium foliaceum*. These species have tertiary plastids of diatom origin and a residual diatom-derived nuclei (Žerdoner Čalasan *et al*., 2018), so it is not inconceivable that a diatom TpSin1 is retained in the organism’s DNA. As a result of this, TpSin1 can probably be considered diatom-specific, and it is the only diatom-specific protein found in the transcriptomes of more than 90% of the MMETSP diatom species analysed.

Nineteen proteins were classified as having a broader phylogenetic distribution with putative homologues outside the diatoms (14 from *T. pseudonana*, four from *C. cryptica*, one from *T. oceanica*), often with BLAST hits in non-diatom Ochrophytes, the Miozoa and the Haptophyta. They were classified as either having a Chromista, or Eukaryote-wide distribution. Twelve of these proteins had characterised domains (e.g. sulfatase, phytase, protease), which were then key drivers of BLAST similarity. It is conceivable that these domains may have unique functions when set in the context of more diatom specific sequence. For example, silica formation in sponges is catalysed by a protein that contains a cysteine protease-like domain (Cha *et al*., 1999).

### Silica-targeting sequences

Although there were no enriched motifs shared by all SSPs, we were interested to see if the SSPs shared sequences that might be responsible for ‘silica targeting’, i.e. directing the proteins into the biosilica cell wall. A TpSil3-derived 13mer peptide containing five lysine residues, was previously shown to be sufficient (along with a signal peptide) for silica targeting of GFP in *T. pseudonana* with high efficiency (Poulsen et al., 2013). Similar so-called penta-lysine clusters matching the regular expression K..K..K.{1,2}K..K are also present in the other previously known silaffins, TpSil1, TpSil2 and TpSil4 (Poulsen et al., 2013). Among the SSPs, 27 (29%) contained at least one penta-lysine cluster, while there were matches in only 1.5% of sequences in the background proteomes excluding SSPs. This raises the question how the other SSPs become targeted to the biosilica cell wall. Poulsen et al. (2013) showed that sequence regions containing at least two K..K motifs with a maximal spacing of 10 residues could mediate silica targeting to some extent (Poulsen et al., 2013). Such regions are present in an additional 40 SSPs, yet 25 SSPs lacked any of the silica targeting sequences, including ten that contain no K..K motifs at all. SSPs devoid of any known silica targeting sequences might rely either on a yet unknown silica targeting mechanism or reach the silica cell wall by forming clusters with one of the SSPs that bear a silica targeting sequence.

### Overview of SSP characteristics

Key outputs of the bioinformatic analyses of the SSPs are summarised in Fig. **6**, which provides an integrated overview how the results of the above analyses relate to one another. Species-specific proteins tend to be disordered over a greater portion of their length compared to other SSPs (mean of 75% vs 45%, Welch two sample t-test, t=7, df=50, p=4.6E-9) and to be of low sequence complexity over a greater portion of their length (mean of 29% vs 15%, t=3.8, df=34, p=0.0006) compared to the other SSPs. Compared to proteins in broader phylogenetic categories, proteins classed as species-specific or Mediophyceae age fall at a higher proportion into one of the eight motif clusters (70% vs. 31%; Fig. **S3**). It is notable that there are few long regions of sequence similarity between the SSPs of Mediophycea or Baciallriophyta age apart from known domains such as RCC1 repeats Instead such SSPs seem to share particular combinations of very short, unconventional motifs within intrinsically disordered protein domains of low amino acid complexity. Currently, there is insufficient understanding of the structure-function correlation in such proteins and their evolution making it impossible to identify ortholog based on sequence analysis. To attempt the identification of putative orthologs, we investigated the location patterns of selected SSPs in the biosilica assuming that othologous proteins should exhibit highly similar location patterns.

**Fig. 6.**
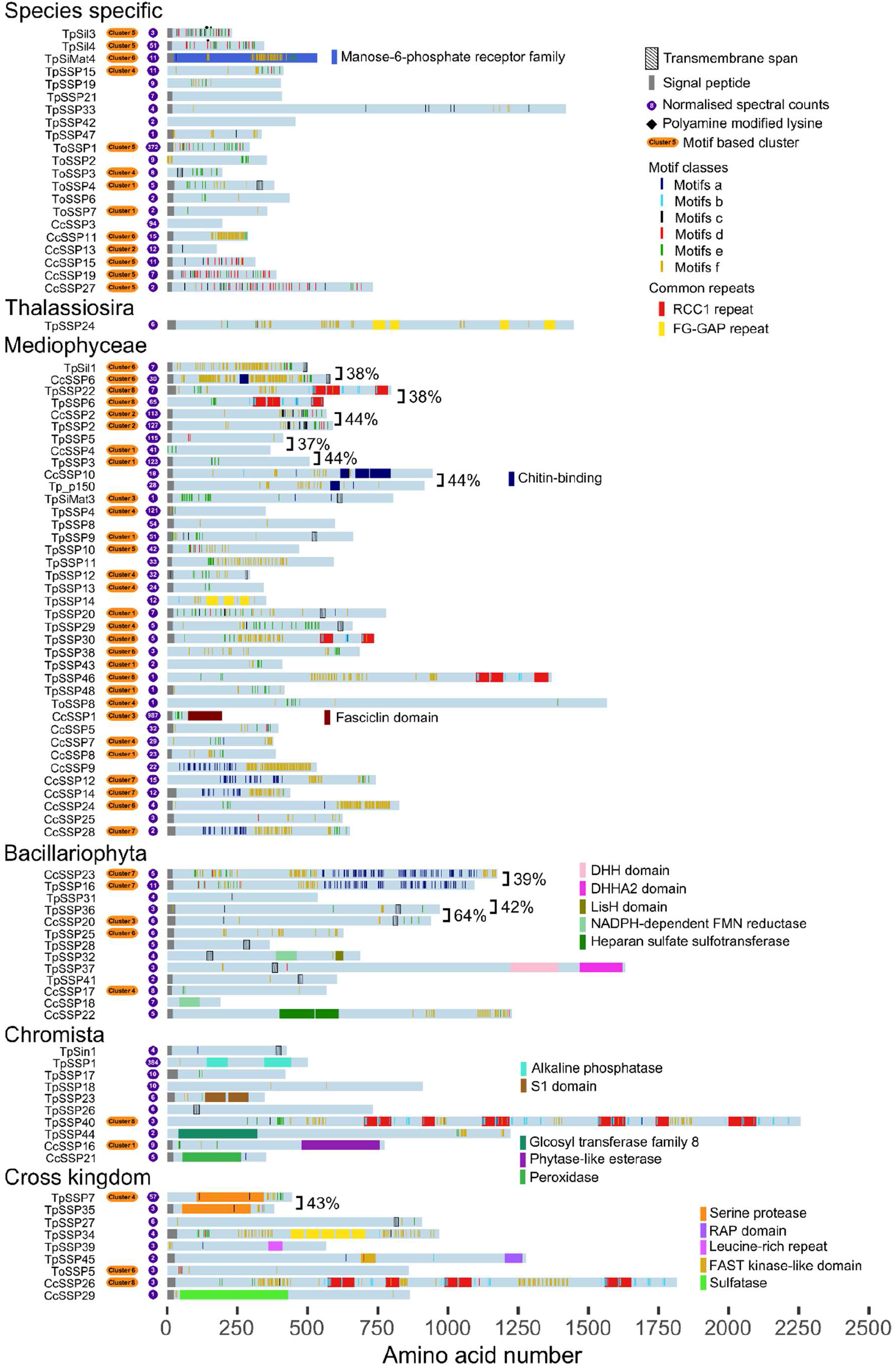
Schematic diagrams summarizing the key features of the SSPs including functional domains, signal peptides and transmembrane domains. The positions of six classes of motif are displayed. The classes and colour coding correspond to Fig. **4**, and cluster membership is indicated. The phylogenetic groupings are derived from the MMETSP data shown in Fig. **5**. The most similar pairs of proteins according to global pairwise sequence alignments are bracketed and their percentage identity shown. The normalised spectral counts (counts per 100 residues) for each protein across all samples is shown to the left of each protein chain with a purple background.

### Localisation of selected SSPs

A selection of SSPs were expressed as GFP fusion proteins in their species of origin and under their native promoters, to validate their silica association in vivo, and to examine their precise positioning with the biosilica. We focused on species-specific, and Mediophyceae specific proteins, because the former might account for species-specific differences in the biosilica structures (e.g. rib patterns), while the latter might have a role in the biogenesis of structural features common to the three species (e.g. fultoportulae, cribrum pores and girdle bands) (see Fig. 1).

In each case, tight association with the silica was tested by extracting the cells with a hot solution of EDTA and SDS, which removes intracellular material and cell wall proteins that are only loosely associated with the silica (Poulsen *et al*. 2013). All except one of the proteins that were GFP-tagged in the present study showed identical localisation patterns in live cells and the purified silica (see below). The GFP-tagging experiments could only be performed with *T. pseudonana* and *C. cryptica*, because transformation experiments using the protocols for both species (Poulsen *et al*., 2006; Kumari *et al*., 2020) did not succeed in obtaining transgenic clones when applied to *T. oceanica*. Schemes showing the position of the GFP tag for each protein are shown in Fig. **S4**.

Given the potential role of silaffins in the morphogenesis of species-specific silica structures (Poulsen *et al*., 2003; Poulsen & Kröger, 2004), we started by examining the locations of some of the *T. pseudonana* silaffins and compared these to the locations of the most similar SSPs that we had identified in *C. cryptica* silica. The closest homolog to silaffin TpSil1 in the *C. cryptica* SSPs is CcSSP6 (36% sequence identity in global alignment, motif-based correlation 0.60). Both proteins are Mediophyceae specific, belong to motif Cluster 6: rich in characteristic ‘type f’ motifs (e.g. PTLSP), but are also in ‘type e’ motifs (e.g. KSGK) (see Fig 4). They also both have a helical region at the extreme C-terminus of the sequence, while TpSil1 has a chitin-binding domain which is absent in CcSSP6. TpSil1-GFP and CcSSP6-GFP were both located exclusively in the valve part of the cell wall, and tightly associated with the silica (Fig. **7a, b**). They were located throughout the valve, but the strongest GFP signal was detected towards the rim. In valve view, CcSSP6-GFP exhibits a sector-type pattern, which is consistent with a localization between the wide ribs of the valve (compare Fig. 7b with Fig. 1h). However this requires further experimental validation.

**Fig. 7.**
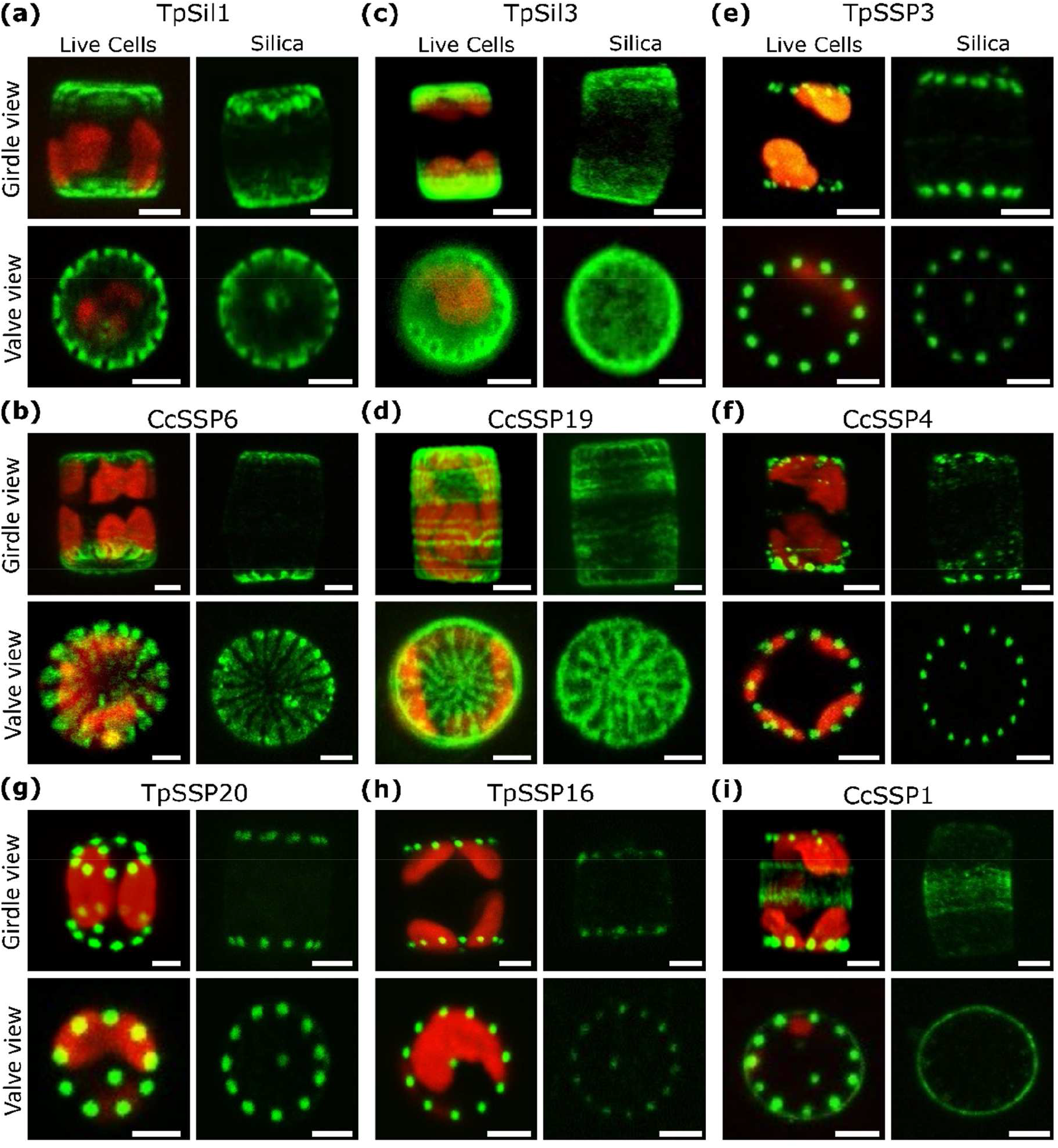
Localization of (a) TpSil1-GFP^int^, (b) CcSSP6^int^-GFP, (c) TpSil3-GFP, (d) CcSSP19-GFP, (e) TpSSP3-GFP, (f) CcSSP4-GFP, (g) TpSSP20-GFP, (h) TpSSP16-GFP and (i) CcSSP1-GFP fusion proteins in *T. pseudonana* and *C. cryptica*. Each protein is expressed in its species of origin. Confocal fluorescence microscopy images are of live cells and isolated biosilica in valve and girdle band orientation. Green: GFP fluorescence, Red: chloroplast autofluorescence. Scale bars: 2 µm.

TpSil3 is structurally the best characterized silaffin of the three diatom species (Poulsen & Kröger, 2004; Sumper et al., 2007; this work), and is located throughout the valves, in the two-girdle bands closest to each valve, and absent from the girdle band in the mid cell region (Fig. **7c**). These observations are consistent with a previous report (Scheffel *et al*., 2011). The most similar *C. cryptica* SSP is CcSSP19. Although they share only 29% sequence identity in global alignments, they have a very high motif-based sequence correlation of 0.89, both belongto Cluster 5, and are both rich in ‘type d’, acidic KxxK motifs such as DAKA.K. Thus, they can be considered to be putative orthologs, despite being classified as species-specific in the phylogenetic analysis (see Fig. 5). Like TpSil3-GFP, CcSSP19-GFP is observed both in the valves and the girdle bands (Fig 7d). However, in contrast to TpSil3-GFP, CcSSP19-GFP appears to be concentrated in radial structures of the valve, which are reminiscent of the pattern and dimensions of the wide ribs (see Fig. **1h**). It also exhibits a striped localization pattern in the girdle bands, with curved regions, which might correspond to non-porous valve regions (Fig **1i**). These data support the idea that both proteins may have similar functions, but with species-specific modulations. Interestingly, it appears that the localizations of CcSSP6 and CcSSP19 may be complementary to one another: the former present predominantly between the wide ribs, and the latter in the wide ribs (compare Fig. 7b and Fig. 7d bottom panels). The more homogeneous distribution of TpSil3-GFP is consistent with the absence of a sector type silica architecture in the *T. pseudonana* valve.

Motif Cluster 1 (rich in ‘type e’ motifs such as KSGK) is the most common cluster among Mediophyceae specific proteins (Fig. **S3**), and did not contain any previously characterised proteins. We chose to examine TpSSP3 and CcSSP4 as representatives of this cluster, which shared 44% identity in a full-length sequence alignment and were identified with high spectral counts (Table S1). Closer inspection of the CcSSP4 gene model revealed that it was truncated, so the non-truncated version (52% identity to TpSSP3), which included a signal peptide, was used for GFP tagging (Fig. **S5**). When expressed as a GFP fusion proteins, TpSSP3 and CcSSP4 were located exclusively in the valves (Fig. **7e, f**). Remarkably, they exhibited an evenly spaced punctate localisation around the circumference of the valve, in a pattern matching the positioning of the fultoportulae (see Fig. 1a, g). We employed correlative fluorescence and electron microscopy (CLEM) imaging to investigate the location of TpSSP3-GFP relative to the fultoportulae in *T. pseudonana*. The result demonstrates that TpSSP3-GFP is indeed precisely co-located with the fultoportulae (Fig. **8a-c**).

**Figure 8.**
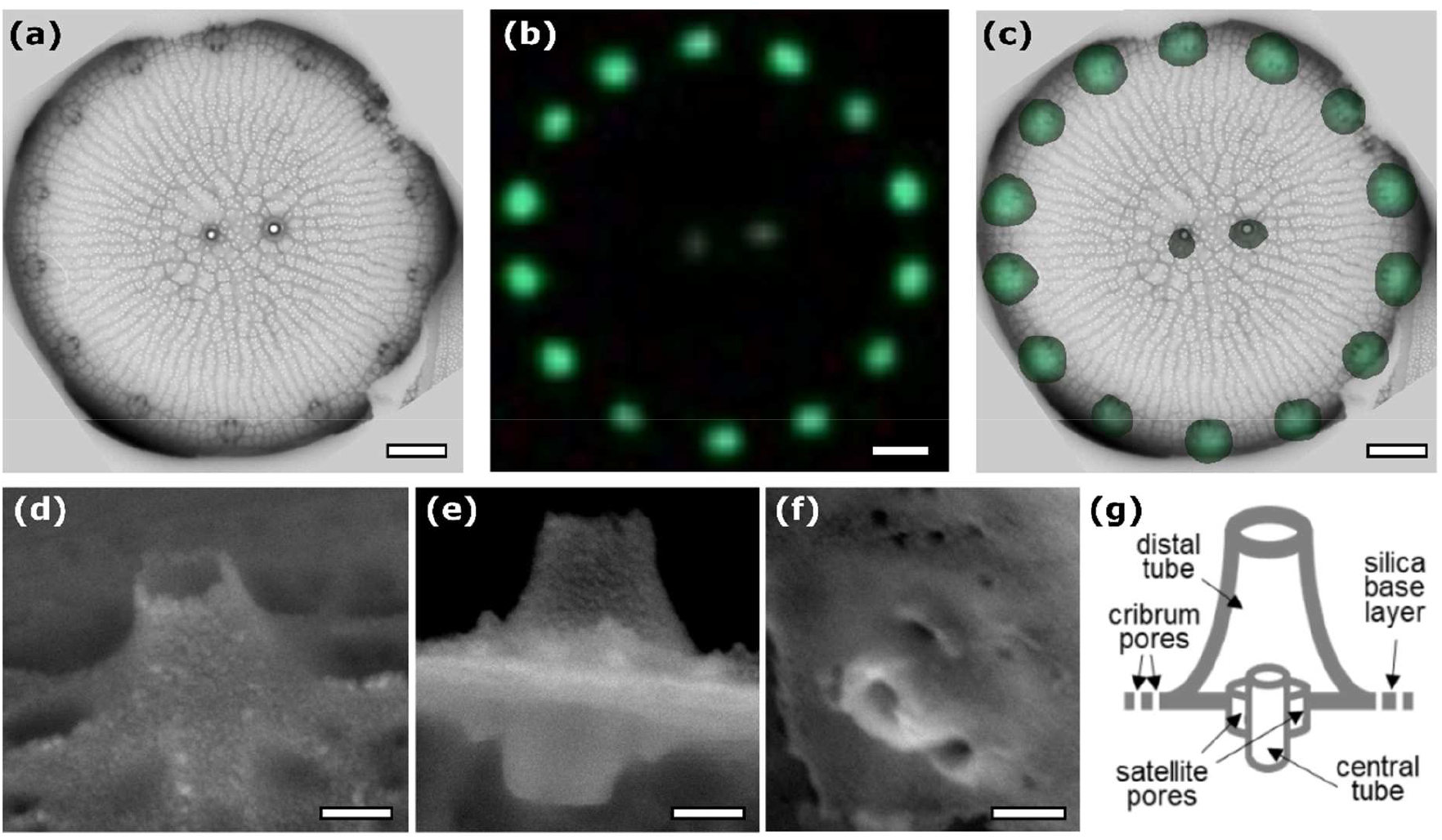
*T. pseudonana* fultoportula structure. (a) TEM image of a *T. pseudonana* valve from a TpSSP3-GFP expressing cell line. (b) GFP fluorescence from the same valve. (c) Overlay of GFP signal and TEM image. Scale bars for a – c are 500 nm. (d-f) Scanning electron microscopy (SEM) images from different individual fultoportulae in *T. pseudonana*. (d) Distal tube in oblique view; (e) view at a valve edge showing the distal tube in the upper half and the central tube with satellite pores on the bottom half of the image; (f) view into the opening of the distal tube showing the central tube and the associated satellite pores; (g) schematic of the fultoportula structure. Scale bars for d – g, 100 nm.

Although also in motif Cluster 1 and well conserved in the Mediophyceae, TpSSP20 differed from the Cluster 1 proteins analysed above (CcSSP3 and CcSSP4) in that it is also enriched in SMSM motifs and contains a single transmembrane domain. Despite these differences it was also located exclusively in the valves and showed a regular, punctate pattern consistent with fultoportulae location (Fig. **7g**).

Motif Cluster 2 also contained only uncharacterised proteins, but all its members have two distinct regions, with N-terminal regions rich in ‘type f’ Ser, Pro and Thr rich motifs, and C-terminal regions rich in ‘type a’ Cys rich motifs such as IGxxSC and GxxACxxN. We chose to localize TpSSP16 as a representative of this group of proteins. It shares 29% sequence identity with CcSSP23. Both these sequences are well conserved across the Mediophyceae, but also have some similar sequences within the Bacillariophyceae, but none in the Coscinodiscophyceae. TpSSP16-GFP also exhibited the characteristic fultoportula localisation pattern (Fig. **7h**). No data could be obtained for CcSSP23-GFP, which failed to display GFP fluorescence in transformant clones.

Given that CcSSP1 lacked similar sequences among *T. pseudonana* SSPs, had a fasciclin domain (that is absent from the TpSSPs) and was identified by more spectral counts than any other *C. cryptica* SSP, we considered CcSSP1 to be a good candidate for a protein which could be responsible for some structural differences between *T. pseudonana* and *C. cryptica* silica. This protein is found in only 6% of Mediophyceae species in the MMETSP data, and belongs to motif Cluster 3, being rich in SxxSSKS motifs as well as ‘type e’ motifs such as KSGK. The GFP fusion protein was observed in both girdle bands and valves. In the girdle bands, CcSSP1-GFP was fairly homogenously distributed in the mid-cell region, and in the valve, it exhibited the familiar fultoportula location pattern (Fig. **7i**). After EDTA/SDS treatment the GFP fluorescence from the fultoportulae was entirely removed, whereas the GFP fluorescence from the girdle bands remained (Fig. **7i**).

## Discussion

The present study vastly increases the number of known silica-associated proteins, providing unique insights into their phylogenetic relationships, sequence characteristics, and possible functions in silica morphogenesis. From the three species, we identified 92 Soluble Silicome Proteins (SSPs), 75 of which had not been previously identified at the protein level. Almost 80% (73 out of 92) of the SSPs appear to be diatom specific, and they are often rich in unusual sequence motifs (Fig. **4**). Sequence similarity among SSPs is generally low: only 10 out of 4186 possible pairs of SSPs displayed a global sequence identity above 35%. It has previously been noted that biomineral-associated proteins tend to be biased in amino acid composition and intrinsically disordered (Evans, 2019). However, such qualitative observations have only rarely been quantitatively analyzed or put into perspective to the whole proteomes of the biomineralizing organisms (Skeffington & Donath, 2020). Here we show that the SSPs differ as a group from their respective background proteomes in (i) being significantly more biased in amino acid composition, (ii) exhibiting significantly lower sequence complexity, and (iii) being predicted to be intrinsically disordered over a larger proportion of their lengths (see Fig. **3**). These trends were more pronounced in species-specific SSPs, although this may simply reflect the difficulties of assigning orthology relationships to low complexity, disordered proteins.

It was very striking, from both our proteomic and SDS-PAGE data, that the soluble silica proteome of *T. oceanica* is quite different from that of *T. pseudonana* and *C. cryptica*. In fact, none of the SSPs from *T. pseudonana* or *C. cryptica* appeared to have homologues in the *T. oceanica* SSP set. This could be interpreted as evidence that the soluble silica proteins are unlikely to be important components of the basal silicification machinery of diatoms, and that their role may be more related to the fine-tuning of the silica structures. However, an alternative explanation is that orthologues (or at least functional homologues) of SSPs do exist in all three species, but they are not readily recognized in sequence alignments and BLAST searches due to the low complexity nature of the sequences. In support of this idea, we found that clustering SSPs by motif content resulted in groups of proteins with members derived from all three species. For example, TpSil3 and TpSil4 belonged to the same cluster as CcSSP19 and ToSSP1, leading us to hypothesise that these proteins may have similar roles in silica formation in each species, driven by the particular physio-chemical properties of the amino acid side chains in the motifs. Indeed, GFP-tagging demonstrated that TpSil3 and CcSSP19 have a very similar distribution within the valve silica. In order to more formally test the functional equivalence of proteins within motif-clusters, knock-out mutants must be generated and characterized. However, the fact that there are often multiple proteins from a species within a cluster indicates that redundancy may make this a challenging approach in practice.

We investigated the intracellular locations of seven newly identified SSPs *in vivo* through GFP-tagging, and each was found to be tightly associated with the biosilica. The result of this spot-check strongly supports the notion that most of SSPs are likely to be genuine silica-associated proteins rather than contaminants. At the same time, we are certain that the set of 92 SSPs does not represent the full complement of silica associated proteins from the three species. In addition to the soluble silicome proteins, the silica of each of the three species is associated with protein based organic matrices that remain insoluble during ammonium fluoride treatment. Only three of the twelve proteins that were previously found in the insoluble organic matrix from *T. pseudonana* were present among the SSPs (TpSil1, TpSiMat3 and TpSiMat4). Consequently, we expect that most of the proteins that constitute the insoluble organic matrices of *C. cryptica* and *T. oceanica* are absent from the SSP data set. To identify the full set of biosilica associated proteins from the three species will require the development of methods to comprehensively characterize the protein complement of the insoluble organic matrices.

An unexpected outcome of the GFP-tagging studies was the identification of five proteins with a fultoportula-specific location, which has never been observed before. Fultoportulae have a much more complex morphology than any of the other structural elements in the cell walls of *Thalassiosira* and *Cyclotella* species. A fultoportula is composed of a central silica tube and two to four satellite pores which each penetrate the silica base layer of the valve (Fig. **8d-g**). At the distal side, the pores are covered by a funnel shaped tube. It is conceivable that the biosynthesis of such an elaborate structure will require a larger number of morphogenetic proteins compared to the comparatively simple patterns of branched, cross-linked ribs and cribrum pores. One of the proteins we found to be fultoportulae-localized, TpSSP20, was previously suggested to be involved in cribrum pore formation based on RNAi experiments targeting the *TpSSP20* gene (Trofimov *et al*., 2019). The resulting mutant phenotype exhibited alterations in the area and density of the cribrum pores in the valve, but no effect on the positioning or morphology of the fultoportulae was reported. Considering the location of TpSSP20, the result from the RNAi work is surprising, yet it is possible that the concentration of the TpSSP20-GFP fusion protein in the cribrum pores was is too low to be detected by fluorescence microscopy. In the RNAi knock-down study there was also no analysis of the degree of TpSSP20 gene silencing. Given the rather mild effect on cribrum pores in the RNAi mutants, it is conceivable that the down-regulation of TpSSP20 achieved was insufficient to affect fultoportula morphogenesis. The generation of knock-out mutants lacking TpSSP20 and the other fultoportula located proteins would provide more definite insight into their role in silica morphogenesis. It would also be interesting to modulate the expression of CcSSP19 and CcSSP6, since their localization suggests that they could have roles in determining the width and spacing of the wide ribs.

Finally, we were interested to see whether any of the SSPs might be part of a basal silicification machinery common to all diatoms. Four groups of homologous proteins had good homologs in more than 80% of diatom species represented in the MMETSP transcriptomes. Given that some of these transcriptomes are likely to be incomplete, and that they are very unlikely to represent the full expression potential of the organism, it is currently reasonable to suppose that these four proteins (TpSSP41, TpSSP28, CcSSP20 (with homologues TpSSP31 and TpSSP36), and TpSin1) are conserved throughout the diatoms. The only one of these proteins to have been previously studied is TpSin1, a knock-out of which displayed changes in silica architecture and reduced the strength of the silica cell wall (Görlich *et al*., 2019). In the future it will be interesting to see if these proteins can be detected in the silica of diatoms from diverse lineages, and to explore their functional roles through reverse genetics.

## Acknowledgements

This work was supported by the Deutsche Forschungsgemeinschaft (DFG) through Research Unit 2038 “Nanomee” (grants KR 1852/8-2 to NK, and SH94/4-2 to ASh) and by a fellowship from the Alexander von Humboldt Stiftung (to AWS). The facility ‘Molecular Analysis - Mass Spectrometry’ at the CMCB was generously supported by grants of the European Regional Development Fund (ERDF/EFRE) (Contract #100232736), the German Federal Ministry of Education and Research (BMBF, 03Z22EB1), and the German Research Foundation (DFG, INST 269/731-1 FUGG). We thank Damian Pawolski for providing the SEM images in Figure 1. We also thank Anke Bormmann, Jennifer Klemm, Jannes Haupstein and Alexander Kotzsch for assistance with cloning of the GFP constructs and diatom transformations.

## Author Contributions

AWS analysed the data, developed bioinformatic methods and wrote the paper. NK and NP conceived the project, designed experiments, analysed data, performed bioinformatics analyses, and wrote the paper. CH, AO, LB, SG designed and performed experiments and analysed data. MG designed proteomic experiments and analysed data.

## References

Alverson AJ, Beszteri B, Julius ML, Theriot EC. 2011. The model marine diatom Thalassiosira pseudonana likely descended from a freshwater ancestor in the genus Cyclotella. BMC Evolutionary Biology 11: Article Number 125.

Benoiston A-S, Ibarbalz FM, Bittner L, Guidi L, Jahn O, Dutkiewicz S, Bowler C. 2017. The evolution of diatoms and their biogeochemical functions. Proceedings of the Royal Society B: Biological Sciences 372: 20160397.

Brembu T, Chauton MS, Winge P, Bones AM, Vadstein O. 2017. Dynamic responses to silicon in Thalasiossira pseudonana - Identification, characterisation and classification of signature genes and their corresponding protein motifs. Scientific Reports 7: Article number: 4865.

Brunner E, Gröger C, Lutz K, Richthammer P, Spinde K, Sumper M. 2009a. Analytical studies of silica biomineralization: Towards an understanding of silica processing by diatoms. Applied Microbiology and Biotechnology 84: 607–616.

Brunner E, Richthammer P, Ehrlich H, Paasch S, Simon P, Ueberlein S, Van Pée KH. 2009b. Chitin-based organic networks: An integral part of cell wall biosilica in the diatom thalassiosira pseudonana. Angewandte Chemie - International Edition 48: 9724–9727.

Buhmann MT, Poulsen N, Klemm J, Kennedy MR, Sherrill CD, Kröger N. 2014. A tyrosine-rich cell surface protein in the diatom Amphora coffeaeformis identified through transcriptome analysis and genetic transformation. PLoS ONE 9: e110369.

Buhmann MT, Schulze B, Förderer A, Schleheck D, Kroth PG. 2016. Bacteria may induce the secretion of mucin-like proteins by the diatom Phaeodactylum tricornutum. Journal of Phycology 52: 463–474.

Campbell KP, MacLennan DH, Jorgensen AO. 1983. Staining of the Ca2+-binding proteins, calsequestrin, calmodulin, troponin C, and S-100, with the cationic carbocyanine dye ‘Stains-all’. Journal of Biological Chemistry 258: 11267–73.

Cha JN, Shimizu K, Zhou Y, Christiansen SC, Chmelka BF, Stucky GD, Morse DE. 1999. Silicatein filaments and subunits from a marine sponge direct the polymerization of silica and silicones in vitro. Proceedings of the National Academy of Sciences of the United States of America 96: 361–365.

Chambers MC, MacLean B, Burke R, Amodei D, Ruderman DL, Neumann S, Gatto L, Fischer B, Pratt B, Egertson J, et al. 2012. A cross-platform toolkit for mass spectrometry and proteomics. Nature Biotechnology 30: 918–920.

Chou MF, Schwartz D. 2011. Biological sequence motif discovery using motif-x. Current Protocols in Bioinformatics 13: 15–24.

Davis AK, Hildebrand M, Palenik B. 2005. A stress-induced protein associated with the girdle band region of the diatom Thalassiosira pseudonana (Bacillariophyta). Journal of Phycology 41: 577–589.

Deutsch EW, Bandeira N, Sharma V, Perez-Riverol Y, Carver JJ, Kundu DJ, García-Seisdedos D, Jarnuczak AF, Hewapathirana S, Pullman BS, et al. 2020. The ProteomeXchange consortium in 2020: Enabling ‘big data’ approaches in proteomics. Nucleic Acids Research 48: D1145–D1152.

Durkin CA, Mock T, Armbrust EV. 2009. Chitin in diatoms and its association with the cell wall. Eukaryotic Cell 8: 1038–1050.

Evans JS. 2019. The Biomineralization Proteome: Protein complexity for a complex bioceramic assembly process. Proteomics 19: 1900036.

Fuentes S, Wikfors GH, Meseck S. 2014. Silicon deficiency induces alkaline phosphatase enzyme activity in cultures of four marine diatoms. Estuaries and Coasts 37: 312–324.

Görlich S, Pawolski D, Zlotnikov I, Kröger N. 2019. Control of biosilica morphology and mechanical performance by the conserved diatom gene Silicanin-1. Communications Biology 2: 245.

Gruber A, Rocap G, Kroth PG, Armbrust EV, Mock T. 2015. Plastid proteome prediction for diatoms and other algae with secondary plastids of the red lineage. Plant Journal 81: 519–528.

Gschloessl B, Guermeur Y, Cock JM. 2008. HECTAR: A method to predict subcellular targeting in heterokonts. BMC Bioinformatics 9: 393.

Harrison PM. 2017. fLPS: Fast discovery of compositional biases for the protein universe. BMC Bioinformatics 18: 476.

Harrison PJ, Waters RE, Taylor FJR. 1980. A broad spectrum artificial sea water medium for coastal and open ocean phytoplankton 1. Journal of Phycology 16: 28–35.

Hecky RE, Mopper K, Kilham P, Degens ET. 1973. The amino acid and sugar composition of diatom cell-walls. Marine Biology 19: 323–331.

Hildebrand M, Frigeri LG, Davis AK. 2007. Synchronized growth of Thalassiosira pseudonana (Bacillariophyceae) provides novel insights into cell-wall synthesis processes in relation to the cell cycle. Journal of Phycology 43: 730–740.

Hildebrand M, Lerch SJL, Shrestha RP. 2018. Understanding diatom cell wall silicification— moving forward. Frontiers in Marine Science 5: 125.

Jones P, Binns D, Chang HY, Fraser M, Li W, McAnulla C, McWilliam H, Maslen J, Mitchell A, Nuka G, et al. 2014. InterProScan 5: Genome-scale protein function classification. Bioinformatics 30: 1236–40.

Kaczmarska I, Beaton M, Benoit AC, Medlin LK. 2005. Molecular phylogeny of selected members of the order Thalassiosirales (Bacillariophyta) and evolution of the fultoportua. Journal of Phycology 42: 121–138.

Kechagia JZ, Ivaska J, Roca-Cusachs P. 2019. Integrins as biomechanical sensors of the microenvironment. Nature Reviews Molecular Cell Biology 20: 457–473.

Keeling PJ, Burki F, Wilcox HM, Allam B, Allen EE, Amaral-Zettler LA, Armbrust EV, Archibald JM, Bharti AK, Bell CJ, et al. 2014. The Marine Microbial Eukaryote Transcriptome Sequencing Project (MMETSP): Illuminating the functional diversity of Eukaryotic life in the oceans through transcriptome sequencing. PLoS Biology 12: e1001889.

Kotzsch A, Gröger P, Pawolski D, Bomans PHH, Sommerdijk NAJM, Schlierf M, Kröger N. 2017. Silicanin-1 is a conserved diatom membrane protein involved in silica biomineralization. BMC Biology 15: 65.

Kotzsch A, Pawolski D, Milentyev A, Shevchenko A, Scheffel A, Poulsen N, Shevchenko A, Kroger N. 2016. Biochemical composition and assembly of biosilica-associated insoluble organic matrices from the diatom Thalassiosira pseudonana. Journal of Biological Chemistry 291: 4982–4997.

Kröger N, Deutzmann R, Sumper M. 1999. Polycationic peptides from diatom biosilica that direct silica nanosphere formation. Science 286: 1129–1132.

Kröger N, Deutzmann R, Bergsdorf C, Sumper M. 2000. Species-specific polyamines from diatoms control silica morphology. Proceedings of the National Academy of Sciences 97: 14133.

Kröger N, Deutzmann R, Sumper M. 2001. Silica-precipitating peptides from diatoms: The chemical structure of silaffin-1A from Cylindrotheca fusiformis. Journal of Biological Chemistry 276: 26066–70.

Kröger N, Lorenz S, Brunner E, Sumper M. 2002. Self-assembly of highly phosphorylated silaffins and their function in biosilica morphogenesis. Science 298: 584–586.

Kröger N, Sumper M. 2004. The molecular basis of diatom biosilica formation. In: Edmund Bäuerlein, ed. Biomineralization: Progress in Biology, Molecular Biology and Application. Wiley Blackwell, 137–158.

Kumari E, Görlich S, Poulsen N, Kröger N. 2020. Genetically Programmed regioselective immobilization of enzymes in biosilica microparticles. Advanced Functional Materials 30: 2000442.

Lachnit M, Buhmann MT, Klemm J, Kröger N, Poulsen N. 2019. Identification of proteins in the adhesive trails of the diatom Amphora coffeaeformis. Philosophical Transactions of the Royal Society B: Biological Sciences 374: 20190196.

Lowenstam HA, Weiner S. 1989. On Biomineralization. New York and London: Oxford University Press.

Marron AO, Ratcliffe S, Wheeler GL, Goldstein RE, King N, Not F, De Vargas C, Richter DJ. 2016. The evolution of silicon transport in eukaryotes. Molecular Biology and Evolution 33: 3226–3248.

Mitchell JG, Seuront L, Doubell MJ, Losic D, Voelcker NH, Seymour J, Lal R. 2013. The role of diatom nanostructures in biasing diffusion to improve uptake in a patchy nutrient environment (S Humphries, Ed.). PLoS ONE 8: e59548.

Mock T, Samanta MP, Iverson V, Berthiaume C, Robison M, Holtermann K, Durkin C, BonDurant SS, Richmond K, Rodesch M, et al. 2008. Whole-genome expression profiling of the marine diatom Thalassiosira pseudonana identifies genes involved in silicon bioprocesses. Proceedings of the National Academy of Sciences of the United States of America 105: 1579–1584.

Nassif N, Livage J. 2011. From diatoms to silica-based biohybrids. Chemical Society Reviews 40: 849–859.

Needleman SB, Wunsch CD. 1970. A general method applicable to the search for similarities in the amino acid sequence of two proteins. Journal of Molecular Biology 48: 443–453.

Nelson DM, Tréguer P, Brzezinski MA, Leynaert A, Quéguiner B. 1995. Production and dissolution of biogenic silica in the ocean: Revised global estimates, comparison with regional data and relationship to biogenic sedimentation. Global Biogeochemical Cycles 9: 359–372.

Nugent T, Jones DT. 2009. Transmembrane protein topology prediction using support vector machines. BMC Bioinformatics 10: Article number: 159.

Osuna-Cruz CM, Bilcke G, Vancaester E, De Decker S, Bones AM, Winge P, Poulsen N, Bulankova P, Verhelst B, Audoor S, et al. 2020. The Seminavis robusta genome provides insights into the evolutionary adaptations of benthic diatoms. Nature Communications 11: 3320.

Pančić M, Torres RR, Almeda R, Kiørboe T. 2019. Silicified cell walls as a defensive trait in diatoms. Proceedings of the Royal Society B: Biological Sciences 286: 20190184.

Pawolski D, Heintze C, Mey I, Steinem C, Kröger N. 2018. Reconstituting the formation of hierarchically porous silica patterns using diatom biomolecules. Journal of Structural Biology 204: 64–74.

Peng K, Radivojac P, Vucetic S, Dunker AK, Obradovic Z. 2006. Length-dependent prediction of protein in intrinsic disorder. BMC Bioinformatics 7: Article number: 208.

Poulsen N, Chesley PM, Kröger N. 2006. Molecular genetic manipulation of the diatom Thalassiosira pseudonana (Bacillariophyceae). Journal of Phycology 42: 1059–1065.

Poulsen N, Kröger N. 2004. Silica morphogenesis by alternative processing of silaffins in the diatom Thalassiosira pseudonana. Journal of Biological Chemistry 279: 42993–42999.

Poulsen N, Scheffel A, Sheppard VC, Chesley PM, Kröger N. 2013. Pentalysine clusters mediate silica targeting of silaffins in Thalassiosira pseudonana. Journal of Biological Chemistry 288: 20100–20109.

Poulsen N, Sumper M, Kröger N. 2003. Biosilica formation in diatoms: Characterization of native silaffin-2 and its role in silica morphogenesis. Proceedings of the National Academy of Sciences of the United States of America 21: 12075–12080.

Qing W, Ni C, Li Z, Li X, Han R, Zhao F, Xu J, Gao X, Wang S. 2019. PureseqTM: efficient and accurate prediction of transmembrane topology from amino acid sequence only. BioRxiv doi.org/10.1101/627307.

Raven JA, Waite AM. 2004. The evolution of silicification in diatoms: Inescapable sinking and sinking as escape? New Phytologist 162: 45–61.

Sapriel G, Quinet M, Heijde M, Jourdren L, Tanty V, Luo G, Le Crom S, Lopez PJ. 2009. Genome-wide transcriptome analyses of silicon metabolism in Phaeodactylum tricornutum reveal the multilevel regulation of silicic acid transporters. PLoS ONE 4: e7458.

Schägger H, von Jagow G. 1987. Tricine-sodium dodecyl sulfate-polyacrylamide gel electrophoresis for the separation of proteins in the range from 1 to 100 kDa. Analytical Biochemistry 166: 368–79.

Scheffel A, Poulsen N, Shian S, Kröger N. 2011. Nanopatterned protein microrings from a diatom that direct silica morphogenesis. Proceedings of the National Academy of Sciences of the United States of America 108: 3175–3180.

Seifert GJ. 2018. Fascinating fasciclins: A surprisingly widespread family of proteins that mediate interactions between the cell exterior and the cell surface. International Journal of Molecular Sciences 19: 1628.

Shrestha RP, Tesson B, Norden-Krichmar T, Federowicz S, Hildebrand M, Allen AE. 2012. Whole transcriptome analysis of the silicon response of the diatom Thalassiosira pseudonana. Bmc Genomics 13: Article number: 499.

Skeffington AW, Donath A. 2020. ProminTools: shedding light on proteins of unknown function in biomineralization with user friendly tools illustrated using mollusc shell matrix protein sequences. PeerJ 8: e9852.

Sumper M, Brunner E. 2008. Silica biomineralisation in diatoms: The model organism Thalassiosira pseudonana. ChemBioChem 9: 1187–94.

Székely GJ, Rizzo ML. 2014. Partial distance correlation with methods for dissimilarities. Annals of Statistics 42: 2382–2412.

Tesson B, Hildebrand M. 2013. Characterization and localization of insoluble organic matrices associated with diatom cell walls: Insight into their roles during cell wall formation. PLoS ONE 8(4): e61675.

Tesson B, Lerch SJL, Hildebrand M. 2017. Characterization of a new protein family associated with the silica deposition vesicle membrane enables genetic manipulation of diatom silica. Scientific Reports 7: 1–13.

Thiele W, Obermaier S, Müller M. 2020. A fasciclin protein is essential for laccase-mediated selective phenol coupling in sporandol biosynthesis. ACS Chemical Biology 15: 844–848.

Thompson LMM, MacLeod RA. 1974. Factors affecting the activity and stability of alkaline phosphatase in a marine pseudomonad. Journal of Bacteriology 117: 813–818.

Traller JC, Cokus SJ, Lopez DA et al. Genome and methylome of the oleaginous diatom Cyclotella cryptica reveal genetic flexibility toward a high lipid phenotype. Biotechnol Biofuels 9, 258 (2016). https://doi.org/10.1186/s13068-016-0670-3

Trofimov AA, Pawlicki AA, Borodinov N, Mandal S, Mathews TJ, Hildebrand M, Ziatdinov MA, Hausladen KA, Urbanowicz PK, Steed CA, et al. 2019. Deep data analytics for genetic engineering of diatoms linking genotype to phenotype via machine learning. npj Computational Materials 5: 67.

Voigt J, Kieß M, Getzlaff R, Wöstemeyer J, Frank R. 2010. Generation of the heterodimeric precursor GP3 of the Chlamydomonas cell wall. Molecular Microbiology.

Volcani BE. 1981. Cell wall formation in diatoms: Morphogenesis and biochemistry. In: Simpson T.L., Volcani B.E., eds. Silicon and Siliceous Structures in Biological Systems. New York, NY: Springer New York.

Vrieling EG, Gieskes WWC, Beelen TPM. 1999. Silicon deposition in diatoms: Control by the pH inside the silicon deposition vesicle. Journal of Phycology 35: 548–559.

Waffenschmidt, S., Woessner, J.P., Beer, K., and Goodenough, U. 1993. Isodityrosine cross-linking mediates insolubilization of cell walls in Chlamydomonas. Plant Cell 5, 809–823.

Wenzl S, Hett R, Richthammer P, Sumper M. 2008. Silacidins: Highly acidic phosphopeptides from diatom shells assist in silica precipitation in vitro. Angewandte Chemie - International Edition 47: 1729–32.

Wieneke R, Bernecker A, Riedel R, Sumper M, Steinem C, Geyer A. 2011. Silica precipitation with synthetic silaffin peptides. Organic and Biomolecular Chemistry 9: 5482–5486.

Willis A, Eason-Hubbard M, Hodson O, Maheswari U, Bowler C, Wetherbee R. 2014. Adhesion molecules from the diatom Phaeodactylum tricornutum (Bacillariophyceae): Genomic identification by amino-acid profiling and in vivo analysis. Journal of Phycology 50: 837–849.

Wootton JC, Federhen S. 1993. Statistics of local complexity in amino acid sequences and sequence databases. Computers and Chemistry 17: 149–163.

Žerdoner Čalasan A, Kretschmann J, Gottschling M. 2018. Absence of co-phylogeny indicates repeated diatom capture in dinophytes hosting a tertiary endosymbiont. Organisms Diversity and Evolution 18: 29–38.

