## Supplementary Information for "Shedding light on biosilica morphogenesis by comparative analysis of the silica-associated proteomes from three diatom species"

Supporting Information

**Fig. S1** Bias in amino acid frequencies for the SSPs of the three species compared to the respective background proteomes. Values above zero indicate enrichment, values below zero indicate depletion.

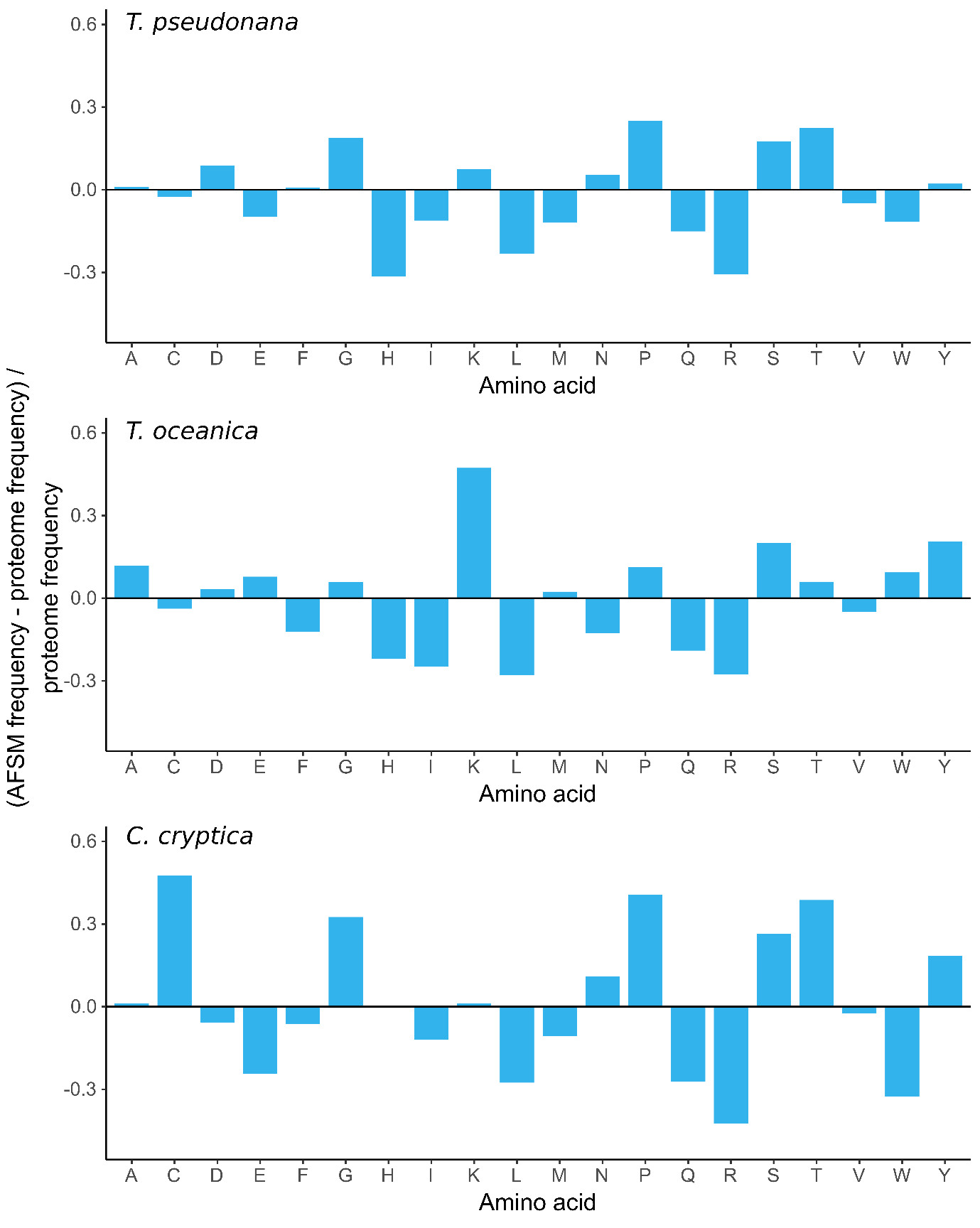

**Fig. S2** Results of all vs. all pairwise blastp comparisons for the protein clusters identified in Fig. 4. The percentage identity of the top HSP for each comparison is indicated by the colour scale and the size of the squares. Query sequences are rows, subject sequences are columns. Note that the nature of the blast comparison means the results are not always exactly symmetrical.

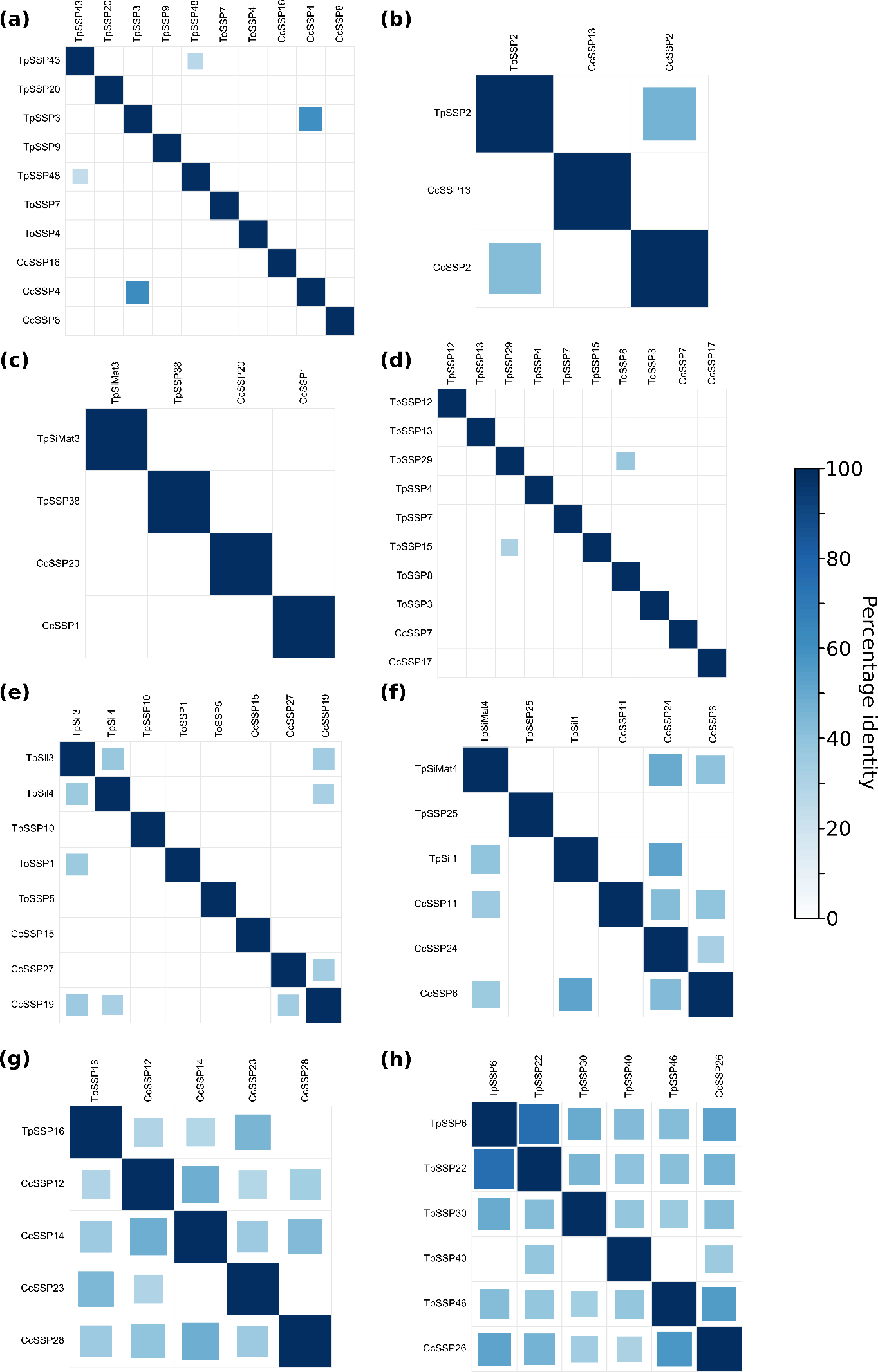

**Fig. S3** Heatmap showing the distribution of protein motif-cluster membership among the phylogenetic categories

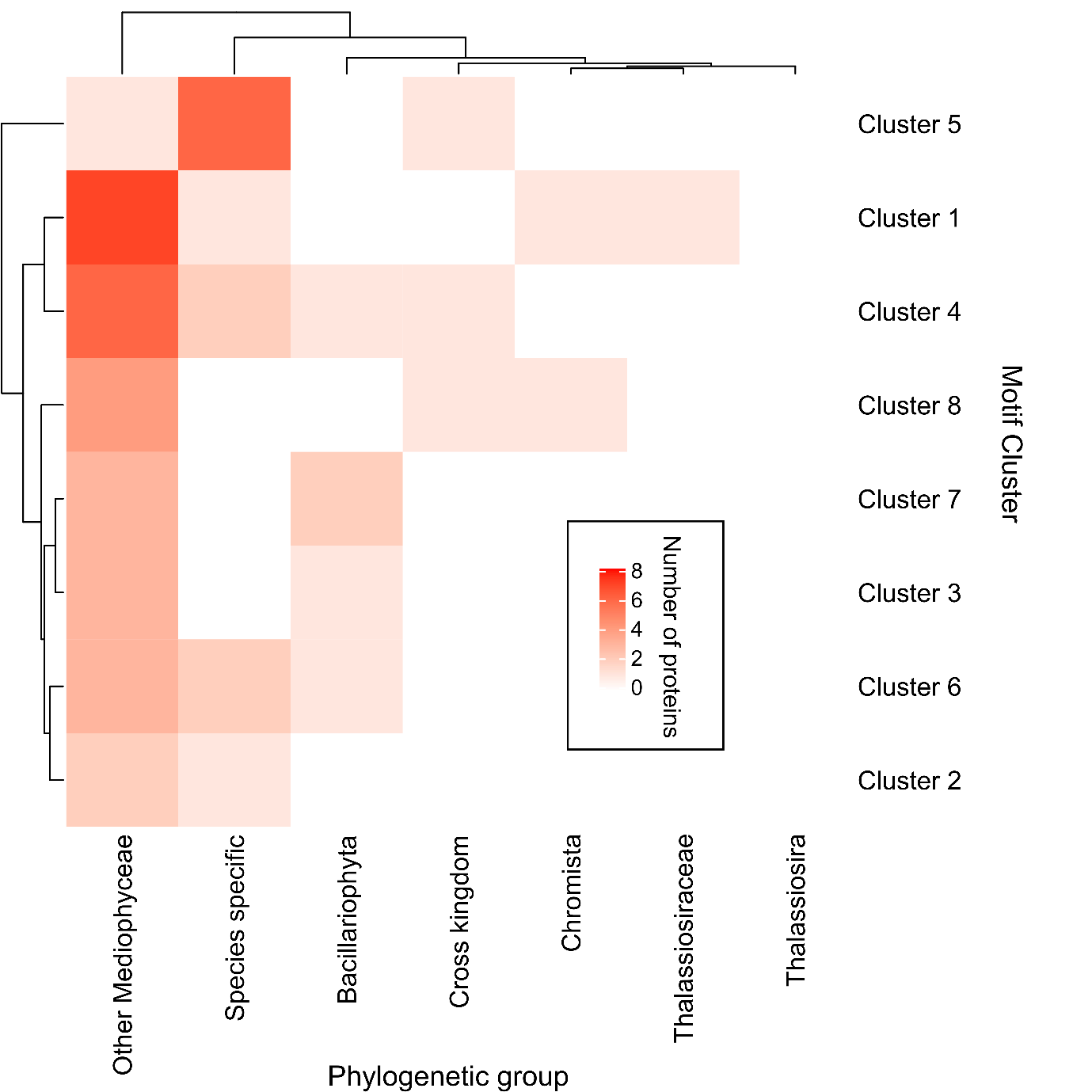

**Fig. S4** Locations of GFP tags in fusion proteins constructed for localization studies

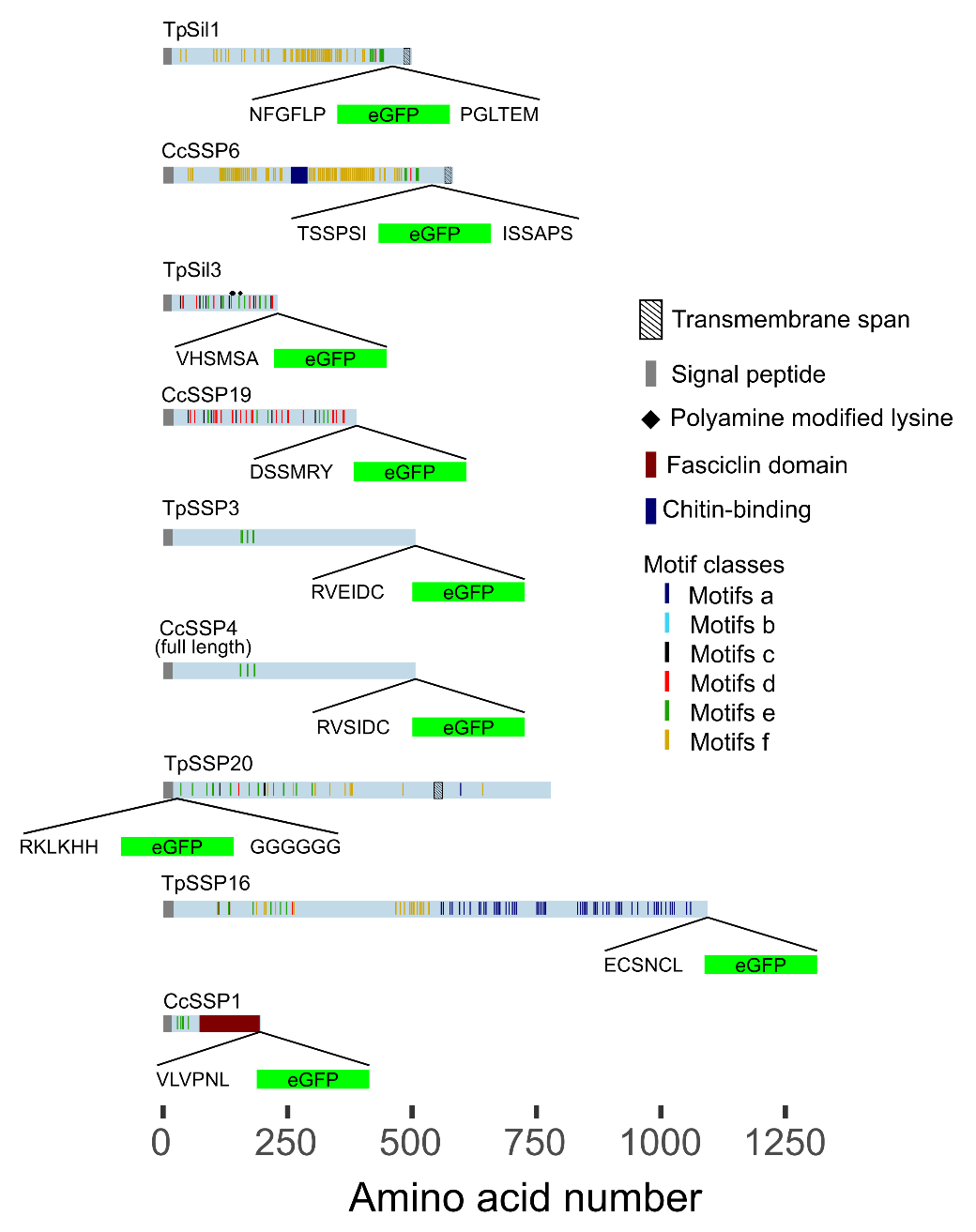

**Fig S5.** The revised gene model of CcSSP4. (a) blast of the CcSSP4 protein sequence (black) against *C. cryptica* genomic contigs revealed two overlapping contigs (green). (b) Predicted cDNA sequence. (c) Predicted protein sequence. The predicted signal peptide in bold and underlined.

(a)

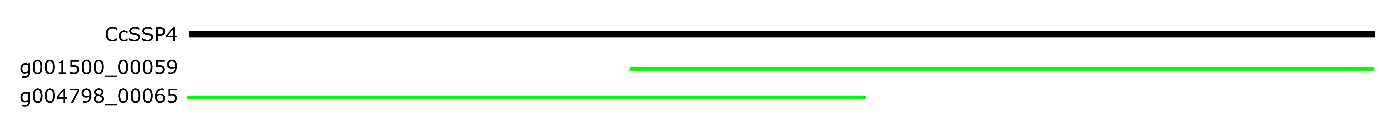

(b)

```
1 ATGAAGACAG TTTCAGTTAT TTTATCCCTC GCCGCAGCTT CAGTTGTAAG AGCGGCCGAA
       61 GATGCAGCTA CTTCTGGAAC AATTCGGGCA TCAAGGAAGG CCACAGACAG CGATGTTAAC
      121 GTCGAATACC CACTTGAAAA GCAGAAGGAG AGGAAAGTTC TCAGCGAGAA TCAACGCAAG
      181 ATGCACCAAG AAGCATTCGA ACGTCATCTC ACGAGCAAGA ACGCGCCGGA GGACGAATCT
      241 GCTGATGTGG TGGATGAGAA GGACGCCAAG AACGCGGACC GCAAGCTATA CAAGAGCAAG
      301 ACGGGTTTGA GCAAGCTCAT GAATGATGAC GGCTATCGCG CCATAAGAAC TGGCGGAAAT
      361 GTTAGACCTC AGGGATGGTT CCAAAATGAT GACATGCCCG GATGGGAAGG TGAGCTGACG
      421 AAAATTTCTT TGTTTTTCAT ATATAGGGAA CTCGAACGCA CACACTTCGT CGGTCTAAAC
      481 TATCCTTCTA GGTGCTGGCA TGCCGGCAAG GATATGGATG ACTGGATGGG AGGTGGTGGA
      541 AAGAGCGGGA AGGATATGGA TGACTGGATG GGCAGTGGCG GAAAGAGCGG TAAGGGCAGC
      601 TCTGGTGGTA ATAACGGATT GTGCCGAGTC ACCGTGACGA ACCTCTCGTT CAAGCAGGCC
      661 TTCAGCAGGA TGTTTGTCAT GTTGCATGAG AAATATGTCA CTCAACACTT CCCTATCTAT
      721 GAGTTTGGAC AGAATCCCTC AAAAGAAATG TGCGATCTTA CACAAAACCT TAACGCGAAC
      781 GGATTGCAAG CCTTTTACAA CGGCCGCCCT GGGGTAGACA GGGCCTGGGT CACATCGAAC
      841 TTTTTTAATT TTGACGACAG CCAACTAGCC ACGACACAAT TCCTTGATGG CGGAGCCACT
      901 TTCAGGTTCT TAGTCCGTCC TATGTCCGGC TACATTTCGC TGGCGGCTGG CTTCCCGTTC
      961 GCTAACGATG GTGGTATAGT GCTCCAAGGA GCTGCAATCT TCAACGGAGC CGAATACTAC
     1021 CTTCCAGCTA TTGATTCTGG ATGTGAAGCA AACTTACAGA CTTGTTGGTC TGTCCCTGCG
     1081 ACCCAAGCAG ACTTTCAAGA ACTAGCACCT CGAGCAGAAT GTGCCGGAGA AAGTCTCTCT
     1141 GATACTAATG ACAACACCTT CGGAGGAGAA AACTTCGTGT CCATCCACCG TGGTATGCAA
     1201 TATTTGAGCG ACAAATCCGA TATCCAAGAA TTGACATTGT TGGAGTGTTC TGATATTTCT
     1261 ATCATCAATG ATGACGATAA CACGAGGTTC GCTAAATACT TCCGTGTTGA GGGCTATGAT
     1321 GATGACCTTC TCCTCTGTGC GTCGGTTGGT GCTGCTGGCC TTTGCAGTTT TAGAGTCGAC
     1381 GACGATTTCC TTGCCTTTGT CAAGTCAAAT GATGATGTTG CCGGAGATGA TGACGTGCTC
     1441 ATCCGCATTG CGAGAAATAG TAATGACTTT AACGACTTCT GCAATCGGAT TGAGGACATC
     1501 AACAAAAATC TCAATTCAGC ATTCAGGGTG CTCGAACCAA TTATCTTCGA TTTTAGAAAT
     1561 CCCATTGCTC GCGTCAGCAT TGACTGTTAA
```

(c)

```
1 MKTVSVILSL AAASVVRAAE DAATSGTIRA SRKATDSDVN VEYPLEKQKE RKVLSENQRK
       61 MHQEAFERHL TSKNAPEDES ADVVDEKDAK NADRKLYKSK TGLSKLMNDD GYRAIRTGGN
      121 VRPQGWFQND DMPGWEGELT KISLFFIYRE LERTHFVGLN YPSRCWHAGK DMDDWMGGGG
      181 KSGKDMDDWM GSGGKSGKGS SGGNNGLCRV TVTNLSFKQA FSRMFVMLHE KYVTQHFPIY
      241 EFGQNPSKEM CDLTQNLNAN GLQAFYNGRP GVDRAWVTSN FFNFDDSQLA TTQFLDGGAT
      301 FRFLVRPMSG YISLAAGFPF ANDGGIVLQG AAIFNGAEYY LPAIDSGCEA NLQTCWSVPA
      361 TQADFQELAP RAECAGESLS DTNDNTFGGE NFVSIHRGMQ YLSDKSDIQE LTLLECSDIS
      421 IINDDDNTRF AKYFRVEGYD DDLLLCASVG AAGLCSFRVD DDFLAFVKSN DDVAGDDDVL
      481 IRIARNSNDF NDFCNRIEDI NKNLNSAFRV LEPIIFDFRN PIARVSIDC*
```

**Table S1** Summary of the characteristics of the SSPs

See separate file

**Table S2** *Thalassiosira pseudonana* SSP genes identified in transcriptomics studies of diatom responses to silicon. Mock *et al.* studied cells under Si and Fe limitation, and whether a gene is up or down regulated in that condition is indicated. Shrestha *et al.* classified genes as ‘silaffin-like response genes’ (SLRGs) if they matched the expression profile of TpSil3 in timecourse after silicon limitation and re-supply of silica. Brembu *et al.* also performed a silica starvation and replenishment experiment. The TpSil2 cluster contained genes that strongly decreased in expression under Si starvation and showed a strong induction after Si replenishment. The CinY1 cluster showed a similar regulation, but of smaller magnitude and the SiMat7-like cluster showed an expression peak 2 – 4 hours after Si replenishment and then expression decreased again.

| SSP ID | Si-limitation  (Mock *et al.*, 2008) | | Silicon starvation  Cluster  (Shrestha *et al.*, 2012) | Silicon starvation and replenishment  clusters and full data set of differentially expressed genes  (Brembu *et al.*, 2017) | | | | Predicted Silaffin-like proteins  (Scheffel et al. 2011) |
| --- | --- | --- | --- | --- | --- | --- | --- | --- |
|  | Si - | Fe - | SLRG | TPSIL2 | CinY1 | SiMat7  -like | Full data | SFLP |
| TpSil3 | X | X | X | X |  |  | X | SFLP87 |
| Tp_p150 | X | X |  |  |  |  | X |  |
| TpSSP3 | X |  | X | X |  |  | X |  |
| TpSSP20 | X |  | X |  | X |  | X | SFLP49 |
| TpSSP16 |  |  | X | X |  |  | X | SFLP15 |
| TpSSP5 |  |  | X |  |  |  | X |  |
| TpSSP11 |  |  | X |  |  |  | X |  |
| TpSSP2 |  |  | X | X |  |  | X | SFLP83 |
| TpSSP12 |  |  | X | X |  |  | X | SFLP56 |
| TpSSP4 |  |  | X |  |  |  | X |  |
| TpSiMat4 |  |  |  |  |  | X | X |  |
| TpSSP39 | X |  | X |  |  |  |  |  |
| TpSSP47 |  |  |  |  |  | X | X |  |
| TpSSP30 |  |  | X | X |  |  | X |  |
| TpSiMat3 |  |  |  |  |  |  | X | SFLP5 |
| TpSSP34 |  |  |  |  | X |  | X | SFLP7 |
| TpSSP29 |  |  |  |  |  |  | X | SFLP20 |
| TpSSP45 |  |  |  |  |  |  | X |  |
| TpSSP35 |  |  |  | X |  |  | X |  |
| TpSSP48 |  |  |  |  |  |  | X | SFLP19 |
| TpSSP14 |  |  |  |  |  |  | X |  |
| TpSSP33 |  |  |  |  |  |  | X | SFLP59 |
| TpSSP21 |  |  |  |  |  |  | X |  |
| TpSSP38 |  |  | X | X |  |  | X |  |
| TpSSP23 |  |  |  |  |  |  | X | SFLP3 |
| TpSSP32 |  |  |  |  |  |  | X |  |
| TpSSP28 |  |  |  |  |  |  | X | SFLP51 |
| TpSin1 |  |  |  |  |  |  | X |  |
| TpSSP40 |  |  |  |  |  |  | X |  |
| TpSSP8 |  |  |  |  |  |  | X |  |
| TpSSP15 |  |  |  |  |  |  | X |  |
| TpSSP43 |  |  |  |  |  |  | X |  |
| TpSSP46 |  |  |  |  |  |  | X |  |
| TpSSP31 |  |  |  |  |  |  | X |  |
| TpSSP26 |  |  |  |  |  |  | X |  |

**Table S3** Results of all vs. all global sequence alignments with EMBOSS Needle.

See separate file. Values are percentage identity.

**Table S4** Motif enrichment values for all SSPs and enriched motifs.

See separate file. Values are fold enrichment relative to the background proteome.

**Table S5** Motif based protein correlation matrix

See separate file

**Table S6** MMETSP transcriptomes used in analyses and taxonomic assignment

See separate file

**Methods S1**

In-solution and in-gel digestion for proteomic analysis by LC-MS/MS were performed according to (Kotzsch *et al.*, 2016) and (Vasilj *et al.*, 2012) respectively, with slight modifications. In brief, for in-solution digests dried extracts were recovered in 10mM ammonium bicarbonate (pH 7.4). Aliquots were digested overnight at 37°C, either with trypsin, chymotrypsin or Asp-N to a final enzyme concentration of 50 ng/µl. Subsequently, the digests were dried and stored at -20°C before recovery in 50µl of 5% formic acid in water, of which 5µl were injected for each LC-MS/MS analysis. For in-gel digests, areas of the gels, either stained with Coomassie G-250 (Sigma) or Stains-All stain (Sigma), were excised and treated for reduction and alkylation of proteins with DTT and iodacetamide. Afterwards proteins were digested with either trypsin, chymotrypsin or Asp-N overnight at 37°C. After peptide extraction , the samples evaoprated to dryness and stored at -20°C. Peptides were recovered in 5% formic acid / water before injection for LC-MS/MS analysis.

LC-MS/MS analyses were performed with a Velos-Orbitrap mass spectrometer or a Q-Exactive HF mass spectrometer (both ThermoScientific, USA/Germany) connected to a nanoflow LC system (Dionex3000 RSLC, ThermoDionex, Germany or Eksigent 2D-LC, Sciex, USA) and operated in data-dependent acquisition (DDA) mode. Spectra from each species were searched against the relevant databases: T. oceanica against Uniprot proteome ID UP000266841, T. pseudonana against Uniprot proteome ID UP000001449 and C. Cryptica against the data base from Taller *et al.* (2016). Each database was supplemented with sequences of common mass spec contaminants and endopeptidases. Peptide and protein identification was performed with Mascot V2.6 Scaff, see below for parameters) and peptide hits were evaluated and aggregated in Scaffold 4.11 (ProteomeSoftware, USA). The proteins were exported from the Scaffold software with the following parameters: protein probability: 99%, peptide probability: 95%, parent ion mass tolerance: 10%, Mascot score cut-off: 20, minimum number of unique peptides per protein: 2. The proteomics data have been deposited at the PRIDE database under the accession PDX026496. Details of instrumentation, parameters and software are listed below.

Instrumentation

LTQ Orbitrap Velos

| Instrument / Parameter | Value | Comments |
| --- | --- | --- |
| LTQ Orbitrap Velos | ThermoScientific, Bremen, Germany | DDA-Mode |
| MS1 |  |  |
| Resolution | R60000 at m/z 400 |  |
| AGC | 1x 10E6 |  |
| Max. Fill Time | 500ms |  |
| Lock Mass | m/z 445.120025 | Dodecamethylcyclohexasiloxane (Schlosser & Volkmer-Engert, 2003) |
| Range | m/z 395-2000 |  |
| Picotip Needle 10µm | 20 µm / 10 µm |  |
| Voltage | 2.7 kV |  |
| MS2 Low Res | Top8 | Low Res. CID (ITMS) |
| AGC | 50000 |  |
| Max. Fill Time | 50ms |  |
| Dynamic Exclusion | 30 s /3 ppm |  |
| Charge states | Unassigned, 1, 5, >5 | (rejected) |
| Threshold | 5E3 |  |
| Isolation Width | 3.0 Da |  |
| Norm. Collision Energy | 35 | (NCE) |
| Activation Q / time | 0.250 / 30 ms |  |
| MS2 High Res |  | High Res. HCD (FTMS) |
| AGC | 5E4 |  |
| Max. Fill Time | 100ms |  |
| Resolution | R7500 at m/z 400 |  |
| Threshold | 5E3 |  |
| Isolation | 3.0 Da |  |
| Fixed 1^st^ Mass | m/z 90 |  |
| Norm. Collision Energy (CE) | 35 | (NCE) |
| Dynamic Exclusion | 30 s /3 ppm |  |
| MS2 High Res |  | High Res. CID (FTMS) |
| AGC | 5E4 |  |
| Max. Fill Time | 100 ms |  |
| Resolution | R7500 at m/z 400 |  |
| Threshold | 5E3 |  |
| Isolation | 3.0 Da |  |
| Norm. Collision Energy (CE) | 35 | (NCE) |
| Dynamic Exclusion | 30 s /3 ppm |  |

Q-Exactive HF

| Instrument / Parameter | Value | Comments |
| --- | --- | --- |
| Q-Exactive HF | ThermoScientific, Bremen, Germany | DDA-Mode |
| MS1 |  |  |
| Resolution | R60000 at m/z 200 |  |
| AGC | 3x 10E6 |  |
| Max. Fill Time | 100ms |  |
| Lock Mass | m/z 445.120025 | Dodecamethylcyclohexasiloxane (Schlosser & Volkmer-Engert, 2003) |
| Range | m/z 375-1700 |  |
| Picotip Needle | 20µm / 10µm |  |
| Voltage | 2.7kV |  |
| MS2 High Res | Top12 | HCD |
| Resolution | R15000 at m/z 200 |  |
| AGC | 1E5 |  |
| Max. Fill Time | 50ms |  |
| Isolation | 1.6 Da |  |
| Fixed 1^st^ Mass | m/z 90 |  |
| Norm. Collision Energy | 23, 25, 27 | (stepped NCE) |
| Threshold | 2.5% / 5E4 |  |
| Charge states | Unassigned, 1, 7,8,>8 | (rejected) |
| Dynamic Exclusion | 15s / 3ppm |  |

Thermo Dionex3000 RSLC

| Instrument / Material | Manufacturer (Supplier) | Comments |
| --- | --- | --- |
| Dionex3000 RSLC | ThermoScientific, Idstein, Germany | Nanoflow System |
| Acclaim PepMap 100 C18,  3 µm, 300 µm x 5 mm, Acclaim PepMap C18 3,  µm, 75 µm x 15 cm | ThermoScientific, Idstein, Germany | Trap-Column Setup Load: 2µl/min Separation: 200nl/min |

Eksigent 2D nanoLC

| Instrument / Material | Manufacturer (Supplier) | Comments |
| --- | --- | --- |
| Eksigent 2D nanoLC | Sciex, Germany | Nanoflow UPLC System |
| Acclaim PepMap 100 C18, 3 µm, 300 µm x 5 mm, Acclaim PepMap C18, 3 µm, 75 µm x 15 cm | ThermoScientific, Idstein, Germany | Trap-Elute Setup Load: 2µl/min Separation: 200nl/min |

Software

| Instrument / Material | Manufacturer (Supplier) | Comments |
| --- | --- | --- |
| MASCOT V2.6 (Perkins *et al.*, 1999) | MatrixScience, London, UK | Protein Identification Software matrixscience.com |
| MSConvert V3.0 (Kessner *et al.*, 2008; Chambers *et al.*, 2012) | Proteowizard, CDN | File Conversion Tool |
| Scaffold V4.11 (Searle, 2010) | Proteome Software, Portland, OR, USA | Protein Identification Statistics and Validation, Visualization  Proteomesoftware.com |

| Mascot Parameters: Low Res. MS2 | Value | Comments |
| --- | --- | --- |
| LTQ Orbitrap-Velos |  |  |
| MS Tolerance | 10 ppm |  |
| Missed Cleavages | 3 / 4 / 4 | Trypsin / Chymotrypsin /AspN |
| Fixed Modifications | (Carbamidomethyl-C) | (Gel slices) |
| Variable Modifications | Acetyl- N-Protein, Oxidation (M) |  |
| MS/MS Tolerance | 0.5 Da |  |
| Instrument | ESI-Trap |  |
| Databases | T. pseudonana (UP000001449) T. oceanica (UP000266841) C.cryptica (custom) Contaminants Enzymes STDs_TAGs | Databases applied according to sample origin. C. cryptica sequences are included in the PRIDE submission |
| Decoy | Yes |  |

| Mascot Parameters: High Res. MS2 | Value | Comments |
| --- | --- | --- |
| LTQ Orbitrap-Velos |  |  |
| MS Tolerance | 10 ppm |  |
| Missed Cleavages | 3 / 4 / 4 | Trypsin / Chymotrypsin /AspN |
| Fixed Modifications | (Carbamidomethyl-C) | (Gel slices) |
| Variable Modifications | Acetyl- N-Protein, Oxidation (M) |  |
| MS/MS Tolerance | 30 mmu |  |
| Instrument | ESI-Trap / ESI-Quad | CID / HCD |
| Databases | T. pseudonana (UP000001449) T. oceanica (UP000266841) C.cryptica (custom) Contaminants Enzymes STDs_TAGs | Databases applied according to sample origin. C. cryptica sequences are included in the PRIDE submission |
| Decoy | Yes |  |

| Mascot Parameters: QE-HF | Value | Comments |
| --- | --- | --- |
| Q-Exactive HF |  |  |
| MS Tolerance | 10 ppm |  |
| Missed Cleavages | 3 / 4 / 4 | Trypsin / Chymotrypsin /AspN |
| Fixed Modifications | (Carbamidomethyl-C) | (Gel slices) |
| Variable Modifications | Acetyl- N-Protein, Oxidation (M) |  |
| MS/MS Tolerance | 30 mmu |  |
| Instrument | ESI-Quad |  |
| Databases | T. pseudonana (UP000001449) T. oceanica (UP000266841) C.cryptica (custom) Contaminants Enzymes STDs_TAGs | Databases applied according to sample origin. C. cryptica sequences are included in the PRIDE submission |
| Decoy | Yes |  |

Chemicals

| Item (Chemicals) | Manufacturer (Supplier) | Order Number |
| --- | --- | --- |
| DTT | Sigma | D-5545 |
| NH_4_HCO_3_ Ammoniumbicarbonate | Sigma | A-6141 |
| IAA | Sigma | I-1149 |
| Water HPLC grade | Merck, Darmstadt, Germany | 1.15333.2500 |
| Acetonitrile HPLC Grade | Merck, Darmstadt, Germany | 1.00029.2500 1.00030.2500 |
| Formic Acid p.a. | Merck, Darmstadt, Germany | 1.00264.0100 |
| Trypsin Gold sequencing grade, (modified Trypsin) | Promega, Walldorf, Germany | V5280 |
| Chymotrypsin Sequencing Grade | Promega, Walldorf, Germany | V1062 |
| Asp-N Sequencing Grade | Promega, Walldorf, Germany | V1621 |

**Methods S2** Clustering proteins based on motif content

Since the aim was to find similarities between SSPs, the clustering analysis was restricted to enriched motifs that were found in at least 5% of the SSPs. The enrichment of each protein for the identified motifs was calculated along with the Distance correlation metric (dcor, using the ‘energy’ package in R) between the motif enrichment profile for each pair of proteins. Proteins were excluded from the analysis if they did not correlate with at least one other protein with a dcor greater than 0.68. The choice of this parameter value was based on manual inspection of clustering results and weighted networks for a range of values, aiming to choose a final value that gave a good representation of the sequence similarity in the data set. Before clustering, four proteins (ToSSP16, CcSSP23, ToSSP12 and CcSSP6) were removed from the analysis since they were only similar to each other because they lacked high enriched motifs. Hierarchical clustering was performed using the Ward.D method with the distance matrix calculated as one minimum the dcor value. An appropriate number of clusters was chosen using the Silhouette method.

**Methods S3** MMETSP blast analysis

Searches using tBLASTn were carried out using custom script mmetspblast3.pl (https://github.com/skeffington/Silicome) which collates the blast results in a taxonomically aware manner. Transcriptomes from the three species that are the focus of this study were excluded from the analysis. We also excluded data from strain simply classed as Thalassiosira without a species name. In preparing the data for Fig. 4, *Tiarina fusus* was also excluded from the results. This is because this transcriptome appears to contain a lot of diatom genes, and the organism is known to predate diatoms, leading us to question the veracity of the blast hits.

**Methods S4** Generating SSP –GFP fusion constructs

Diagrams of all fusion proteins generated are provided in Fig S3.

*Genomic DNA isolation*

Genomic DNA was isolated from *T. pseudonana* and *C. cryptica* according to a previously published method (Poulsen *et al.*, 2006).

*Fusion gene TpSil1-GFPint*

For expression of the tpSil1-GFP fusion protein the 5`-end of the tpSil1 gene together with 980 bp of the promoter and 588bp of the terminator region were amplified from gDNA using the sense primer 5`-CAA GGC AAA AGA GAC AGA-3` and the antisense primer 5`-GAA GGA GAG AGG AAG ACA-3`. The 3997 bp PCR product was inserted into the pJET1.2/blunt Cloning Vector, resulting in plasmid Pjet1.2/tpSil1. The enhanced green fluorescent protein (eGFP) gene was amplified by PCR and then introduced blunt into the SmaI site of Pjet1.2/tpSil1, generating an internal GFP fusion protein as described previously (Poulsen *et al.*, 2013).

*Fusion gene CcSSP6-GFPint*

To construct ccSSP6-GFP fusion protein, a BLAST (Basic Local Alignment Search Tool) was performed comparing the ccSSP6 protein sequence against the six-frame translated C. cryptica genome data using the CLC workbench software. This alignment was manually analyzed in order to identify missed annotations due to gaps in the genome assembly. This verification was done to all proteins that were not previously characterized. After that, the 5`-end of the CcSSP6 gene together with 699 bp of the promoter and 255bp of the terminator region were amplified from gDNA using the sense primer 5`-TAA GCA GGA TCC GCG AGA AGG AAT CTA AG-3` and the antisense primer 5`-TGC TTA GGA TCC ATA CAA ATG ATA CGA CGC CA-3`. The resulting PCR product was digested with BamHI (presented in bold in the oligonucleotides sequences) and inserted into pCcfcpNat (Ekta`s paper) plasmid, which was linearized with the same restriction enzyme, generating then the construct pCcfcpNat/ccSSP11. The eGFP gene was amplified by PCR using the sense primer 5`-AAT GGT GAG CAA GGG CGA-3` and the antisense primer 5`-TGG ACG AGC TGT ACA AGA T-3` and introduced blunt into the EcoRV site of pCcfcpNat/ccSSP6. The final plasmid, pCcfcpNat/ccSSP6-GFPint, resulted in a GFP fusion protein with an internal GFP located after the amino acid position 432 of CcSSP6.

*Fusion gene TpSSP3-GFP*

The TpSSP3 gene together with 1001 bp of the promoter was PCR amplified from gDNA using the sense primer 5’-GCC CGG GGG ATC CAC TAG TTA AGA ACA TGT TAC TAA CCT G-3’ and the antisense primer 5’-TGC TCA CCA TAC AAT CGA TTT CAA CGC G-3; the eGFP gene was amplified using the sense primer 5’- AAT CGA TTG TAT GGT GAG CAA GGG CGA G-3’ and the antisense 5’- TTT GTC ATA TTT ACT TGT ACA GCT CGT CCA TG-3’. The tpSSP3 gene terminator (523 bp) was amplified from gDNA using the sense primer 5’-GTA CAA GTA AAT ATG ACA AAT TCC TTG CGC TG-3’ and the antisense primer 5’-CCA CCG CGG TGG CGG CCG CTA AAC TCA CTC GCT CGC CAC-3’. The pBluescript SK(+) plasmid was linearized with XbaI, and was assembled with the PCR fragments in a NEBuilder® HiFi DNA assembly reaction (NEB), thereby generating the construct pBluescript/tpSSP3-GFP with a C-terminal GFP tag. Since the final construct does not have the selectable marker for T. pseudonana, a co-transformation was performed with pTpfcpNat plasmid.

*Fusion gene CcSSP4-GFP*

The *CcSSP4* gene together with 1154 bp of the promoter was PCR amplified from gDNA using the sense primer 5’-ATT CGA GCT CGG TAC CCG GGC ATA TCA GGT TTG TCG GGT G-3’ and the antisense primer 5’-TGC TCA CCA TAC AGT CAA TGC TGA CGC G-3; the eGFP gene was amplified using the sense primer 5’-CAT TGA CTG TAT GGT GAG CAA GGG CGA G-3’ and the antisense 5’-CTT CGT CTT CTT ACT TGT ACA GCT CGT CCA TG-3’. The ccSSP9 gene terminator (394 bp) was amplified from gDNA using the sense primer 5’-GTA CAA GTA AGA AGA CGA AGA AAG TGT GC -3’ and the antisense primer 5’-GCC CAG CGT GGC ATT TCT CGG GAT GGG AAG CAT GTA GTG-3’. The pCcfcpNat plasmid was linearized with BamHI, and was assembled with the PCR fragments in a NEBuilder® HiFi DNA assembly reaction (NEB), thereby generating the final plasmid pCcfcpNat/ccSSP9-GFP with a C-terminal GFP tag.

*Fusion gene TpSSP16-GFP*

The *TpSSP16* gene together with 1000 bp of the promoter was PCR amplified from gDNA using the sense primer 5’-TCC CGG GGG ATC CAC TAG TTA GTC CGA CAT GTG GAG TG-3’ and the antisense primer 5’-TGC TCA CCA TCA AGC AGT TGC TAC ATT CAC -3; the eGFP gene was amplified using the sense primer 5’- CAA CTG CTT GAT GGT GAG CAA GGG CGA G-3’ and the antisense 5’- ATC TCT TTC CTT ACT TGT ACA GCT CGT CCA TG-3’. The tpSSP5 gene terminator (500 bp) was amplified from gDNA using the sense primer 5’-GTA CAA GTA AGG AAA GAG ATT GGG GGG G-3’ and the antisense primer 5’- CGC GGT GGC GGC CGG CCG CTG ACG TTG GTC AGG TCT TAG-3’. The pBluescript SK(+) plasmid was linearized with XbaI, and was assembled with the PCR fragments in a NEBuilder® HiFi DNA assembly reaction (NEB), thereby generating the construct pBluescript/tpSSP5-GFP with a C-terminal GFP tag. Since the final construct does not have the selectable marker for *T. pseudonana*, a co-transformation was performed with pTpfcpNat plasmid.

*Fusion gene TpSSP20-GFP*

For the N-terminal GFP-tagging of TpSSP20, the GFP gene was inserted after amino acid position 26, downstream of the RKL motif of TpSSP20. The region of the *TpSSP20* gene upstream of the RKL motif, including the promoter region was amplified from genomic DNA using the sense primer 5’- CGT ACC GGG ACC CCC CTC GAG TAG GCC GCC GTC GTC TTT G-3’ and the antisense primer 5’-CCT TGC TCA CCA TTC CGC CGG TAC CAT GAT GTT TGA GCT TTC GTA C-3’. The PCR product was introduced into the ApaI and KpnI sites of pPSin1-Sin1Part1-GFPN-Sin1Part2-TSin1/fcpNAT (Kotzsch *et al.*, 2017) by NEBuilder® HiFi DNA assembly reaction (NEB) resulting in the vector p TpSSP1-SM01Part1-GFPN-Sin1Part2-TSin1/fcpNAT. The TpSSP20 coding region downstream of the RKL motif including the terminator region was amplified from genomic DNA using the sense primer 5’- AGC TGT ACA AGG GAG CGG CCG CTG CGG GTG GAG GAG GAG GCA AGT C-3’ and the antisense primer 5’- GGA AAA AGC GCA AGC TGG GAT GCA TAG TGC AAA CTA GTA GAC CAG-3’. The PCR product was introduced into the NotI and NsiI sites of tpSSP1-SM01Part1-GFPN-Sin1Part2-TSin1/fcpNAT by NEBuilder® HiFi DNA assembly reaction (NEB), generating the final construct.

*Fusion gene CcSSP1-GFP*

The *CcSSP1* gene together with 734 bp of the promoter was PCR amplified from gDNA using the sense primer 5’-CGG CCA GTG AAT TCG AGC TCT TCT CTC CTA AGT AAG TTA TTT TTA CG-3’ and the antisense primer 5’-GCT CAC CAT ACC GCT AGC TCC CAA GTT AGG AAC AAG AAC AGT ATC-3; the eGFP gene was amplified using the sense primer 5’-CTA ACT TGG GAG CTA GCG GTA TGG TGA GCA AGG GCG AG-3’ and the antisense 5’- GTT TCA AGT CTG CCATGG TTA TTT GTA CAG CTC GTC CAT G-3’. The *CcSSP1* gene terminator (542 bp) was amplified from gDNA using the sense primer 5’- GAG CTG TAC AAA TAA CCA TGG CAG ACT TGA AAC GCA GGA ATG G -3’ and the antisense primer 5’-AGC GTG GCA TTT CTC GGA TCC GAA AAA GAA TGT TGC ATT AAT GGT ATG G-3’. The pCcfcpNat plasmid was linearized with BamHI, and was assembled with the PCR fragments in a NEBuilder® HiFi DNA assembly reaction (NEB), thereby generating the final plasmid pCcfcpNat/ CcSSP1-GFP with a C-terminal GFP tag.

*Fusion gene CcSSP19-GFP*

For C-terminal GFP-tagging of CcSSP19, the *CcSSP19* terminator region (522bp downstream of the stop codon) was amplified from genomic DNA using the sense primer 5’- TTC AGC GGC CGC ATA ACT GGA AAA GTG CTT GAA G-3’ and the antisense primer 5’- TGC CGT TAA CGA CAT CCG ACT ATA AGG AGG AT-3’ (NotI site underlined and HpaI site in bold). The resulting PCR product was digested with NotI and HpaI and introduced into the NotI and HpaI sites of pTpNR-GFPHpaI/fcpNat(-NotI) (Kotzsch *et al.*, 2016). The promoter region (579 bp upstream of the start codon) and protein coding region of CcSSP19 were amplified from genomic DNA using the sense primer 5’- TAC CGG GCC CGC AAA GAT CAG GAC-3’ and the antisense primer 5’- AAT GGT ACC ATA CCG CAT TGA GCT GTC TCC-3’ (ApaI site underlined, KpnI site in bold) and introduced into the ApaI and KpnI sites of pPnr-ccSSP2/fcpNAT(-NotI) generating the final expression plasmid PccSSP2-ccSSP2-GFPC-TccSSP2/fcpNAT(-NotI). In this plasmid ccSSP19 is expressed with a C-terminal GFP-tag.

**Methods S5:** Confocal fluorescence microscopy imaging

The imaging of live cells and silica cell walls of transformants expressing GFP fusion proteins was performed using a Zeiss LSM 780 confocal equipped an argon laser (488 nm) and a 32-channel GaAsP spectral detector (acquisition between 473 and 668 nm). Two channels were acquired to separately monitor the GFP fluorescence (emission at 491–535 nm) and chloroplast fluorescence (emission at 654– 693 nm). Images were analysed using the ZEN 3.1 blue edition software (Zeiss).

To determine whether the GFP fusion proteins were incorporated into the silica, cells were incubated at 55°C for 1 h with a buffer containing 1% (w/v) SDS, 0.1 M EDTA (pH 8.0), and 1 mM PMSF. The extracted cell walls were washed three times with water, once with acetone, and again three times with water before imaging.

**References**

**Brembu T, Chauton MS, Winge P, Bones AM, Vadstein O**. **2017**. Dynamic responses to silicon in Thalasiossira pseudonana - Identification, characterisation and classification of signature genes and their corresponding protein motifs. *Scientific Reports*.

**Chambers MC, MacLean B, Burke R, Amodei D, Ruderman DL, Neumann S, Gatto L, Fischer B, Pratt B, Egertson J, *et al.*** **2012**. A cross-platform toolkit for mass spectrometry and proteomics. *Nature Biotechnology* **30**: 918–920.

**Kessner D, Chambers M, Burke R, Agus D, Mallick P**. **2008**. ProteoWizard: Open source software for rapid proteomics tools development. *Bioinformatics* **24**: 2534–2536.

**Kotzsch A, Gröger P, Pawolski D, Bomans PHH, Sommerdijk NAJM, Schlierf M, Kröger N**. **2017**. Silicanin-1 is a conserved diatom membrane protein involved in silica biomineralization. *BMC Biology* **15**: 65.

**Kotzsch A, Pawolski D, Milentyev A, Shevchenko A, Scheffel A, Poulsen N, Shevchenko A, Kroger N**. **2016**. Biochemical composition and assembly of biosilica-associated insoluble organic matrices from the diatom Thalassiosira pseudonana. *Journal of Biological Chemistry* **291**: 4982–4997.

**Mock T, Samanta MP, Iverson V, Berthiaume C, Robison M, Holtermann K, Durkin C, BonDurant SS, Richmond K, Rodesch M, *et al.*** **2008**. Whole-genome expression profiling of the marine diatom Thalassiosira pseudonana identifies genes involved in silicon bioprocesses. *Proceedings of the National Academy of Sciences of the United States of America* **105**: 1579–1584.

**Perkins DN, Pappin DJC, Creasy DM, Cottrell JS**. **1999**. Probability-based protein identification by searching sequence databases using mass spectrometry data. *Electrophoresis* **20**: 3551–3567.

**Poulsen N, Chesley PM, Kröger N**. **2006**. Molecular genetic manipulation of the diatom Thalassiosira pseudonana (Bacillariophyceae). *Journal of Phycology* **42**: 1059–1065.

**Poulsen N, Scheffel A, Sheppard VC, Chesley PM, Krog̈er N**. **2013**. Pentalysine clusters mediate silica targeting of silaffins in Thalassiosira pseudonana. *Journal of Biological Chemistry* **288**: 20100–20109.

**Schlosser A, Volkmer-Engert R**. **2003**. Volatile polydimethylcyclosiloxanes in the ambient laboratory air identified as source of extreme background signals in nanoelectrospray mass spectrometry. *Journal of Mass Spectrometry* **38**: 523–525.

**Searle BC**. **2010**. Scaffold: A bioinformatic tool for validating MS/MS-based proteomic studies. *Proteomics* **10**: 1265–1269.

**Shrestha RP, Tesson B, Norden-Krichmar T, Federowicz S, Hildebrand M, Allen AE**. **2012**. Whole transcriptome analysis of the silicon response of the diatom Thalassiosira pseudonana. *Bmc Genomics* **13**.

**Traller JC, Cokus SJ, Lopez DA, Gaidarenko O, Smith SR, McCrow JP, Gallaher SD, Podell S, Thompson M, Cook O, *et al.*** **2016**. Genome and methylome of the oleaginous diatom Cyclotella cryptica reveal genetic flexibility toward a high lipid phenotype. *Biotechnology for Biofuels* 9:258  https://doi.org/10.1186/s13068-016-0670-3

**Vasilj A, Gentzel M, Ueberham E, Gebhardt R, Shevchenko A**. **2012**. Tissue Proteomics by One-Dimensional Gel Electrophoresis Combined with Label-Free Protein Quantification. *Journal of Proteome Research* **11**.
